# Nuclear Pore Complex Acetylation Regulates mRNA Export and Cell Cycle Commitment in Budding Yeast

**DOI:** 10.1101/2021.09.01.458533

**Authors:** Mercè Gomar-Alba, Vasilisa Pozharskaia, Celia Schaal, Arun Kumar, Basile Jacquel, Gilles Charvin, J. Carlos Igual, Manuel Mendoza

## Abstract

Nuclear pore complexes (NPCs) mediate communication between the nucleus and the cytoplasm and regulate gene expression by interacting with transcription and mRNA export factors. Lysine acetyl-transferases (KATs) promote transcription through acetylation of chromatin-associated proteins. We find that Esa1, the KAT subunit of the yeast NuA4 complex, also acetylates the nuclear pore basket component Nup60 to promote mRNA export. Acetylation of Nup60 recruits to the nuclear basket the mRNA export factor Sac3, the scaffolding subunit of the Transcription and Export 2 (TREX-2) complex. Esa1-dependent nuclear export of mRNAs promotes entry into S phase, and is inhibited by the Hos3 deacetylase in G1 daughter cells to restrain their premature commitment to a new cell division cycle. This mechanism also inhibits expression of the nutrient-regulated *GAL1* gene specifically in daughter cells. These results reveal how acetylation contributes to the functional plasticity of NPCs in specific cell types, and demonstrate how the evolutionarily conserved NuA4 complex regulates gene expression dually at the level of transcription and mRNA export, by modifying the nucleoplasmic entrance to nuclear pores.

## Introduction

Nuclear pores are macromolecular assemblies composed of approximately 30 different nucleoporins that form a channel across the nuclear envelope (Knockenhauer and Schwartz, 2016; Hampoelz et al., 2019; Raices and D’Angelo, 2021). The central channel mediates communication between the nucleus and cytoplasm. Other NPC sub-structures include the cytoplasmic filaments and the nuclear basket, associated with the cytoplasmic and nuclear sides of the central channel, respectively. The nuclear basket regulates gene expression through interactions with active genes (Casolari et al., 2004; Cabal et al., 2006; Light et al., 2010; Brickner et al., 2019) and with regulators of transcription (Texari et al., 2013; Schneider et al., 2015) and mRNA export (Fischer et al., 2002; Dieppois et al., 2006). These and other studies have suggested that NPCs can act as regulatory sites for the coordination of transcript elongation, processing and export (Sood and Brickner, 2014; Ibarra and Hetzer, 2015).

The nuclear basket also recruits lysine acetyltransferases (KATs) and deacetylases (KDACs). Acetylation of histones is tightly associated with active transcription, and KATs and KDACs, often residing in large multiprotein complexes, are thought to target nucleosomes to regulate transcriptional activity (Sterner and Berger, 2000; Lee and Workman, 2007). In addition, acetylation of non-histone proteins is common in eukaryotes and has been implicated in a variety of biological processes in addition to transcription, such as DNA damage repair, cell division and signal transduction (Kaluarachchi Duffy et al., 2012; Narita et al., 2019). Among the best-characterised yeast KATs are Esa1 and Gcn5, contained in the NuA4 and SAGA complexes, respectively. Esa1 (known as Kat5 or Tip60 in mammals) is the only essential KAT in budding yeast and is involved in DNA transcription and repair (Allard et al., 1999; Clarke et al., 1999; Doyon and Côté, 2004; Bruzzone et al., 2018). Gcn5 is not essential but plays a role in the transcription of most yeast genes (Baptista et al., 2017; Bruzzone et al., 2018). KATs and KDACs known to associate with NPCs include Gcn5 and the type II deacetylases Hos3 (in yeast) and HDAC4 (in mammals) (Cabal et al., 2006; Kurshakova et al., 2007; Kehat et al., 2011; Kumar et al., 2018). Despite their presence at NPCs, how KATs and KDACs act to regulate gene expression at these sites is poorly understood.

Regulatory principles of the G1/S transition (known as Start in yeast) are evolutionarily conserved: activation of cyclin-dependent kinase (CDK) drives the transcription of hundreds of genes involved in the start of S-phase (Bertoli et al., 2013). Indeed, defects in G1/S control are tightly associated with oncogenesis. For example, pRB is a repressor of the G1/S transition thought to be functionally inactivated in most tumor cells (Frolov and Dyson, 2004). In both yeast and animal cells, inhibition of premature G1/S transition involves the targeting of KDACs to chromatin, generating an environment that is unfavourable for transcription (Frolov and Dyson, 2004; Huang et al., 2009; Takahata et al., 2009; Wang et al., 2009). We recently discovered that in budding yeast, NPC acetylation regulates the G1/S transition (Kumar et al., 2018; Gomar-Alba and Mendoza, 2019). The KDAC Hos3 is cytoplasmic during interphase, but associates with the yeast division site (the mother-bud neck) in mitosis and then binds to daughter-cell NPCs as they traverse the bud neck during anaphase, leading to Hos3 association with the nuclear basket specifically in daughter cells (**Figure 1A**). Hos3-dependent deacetylation of central pore channel nucleoporins in daughter cells enhances nuclear accumulation of the main Start inhibitor (the transcriptional repressor Whi5, functional analog of pRB). In addition, deacetylation of the nuclear basket nucleoporin Nup60 has relatively minor effects on Whi5 nuclear accumulation but is associated with the perinuclear tethering and silencing of a key cell cycle control gene (encoding the G1/S cyclin Cln2, homologue of mammalian Cyclin E) (Kumar et al., 2018). Thus, acetylation of specific nucleoporins promotes different aspects of nuclear pore function necessary for S phase entry, and their deacetylation in daughter cells reinforces cell size control mechanisms that prevent premature S phase in small daughters (Turner et al., 2012). However, the identity of the KAT(s) targeting the NPC for acetylation is unknown, and the molecular mechanism by which NPC acetylation status affects S phase entry remains unclear.

**Figure 1:**
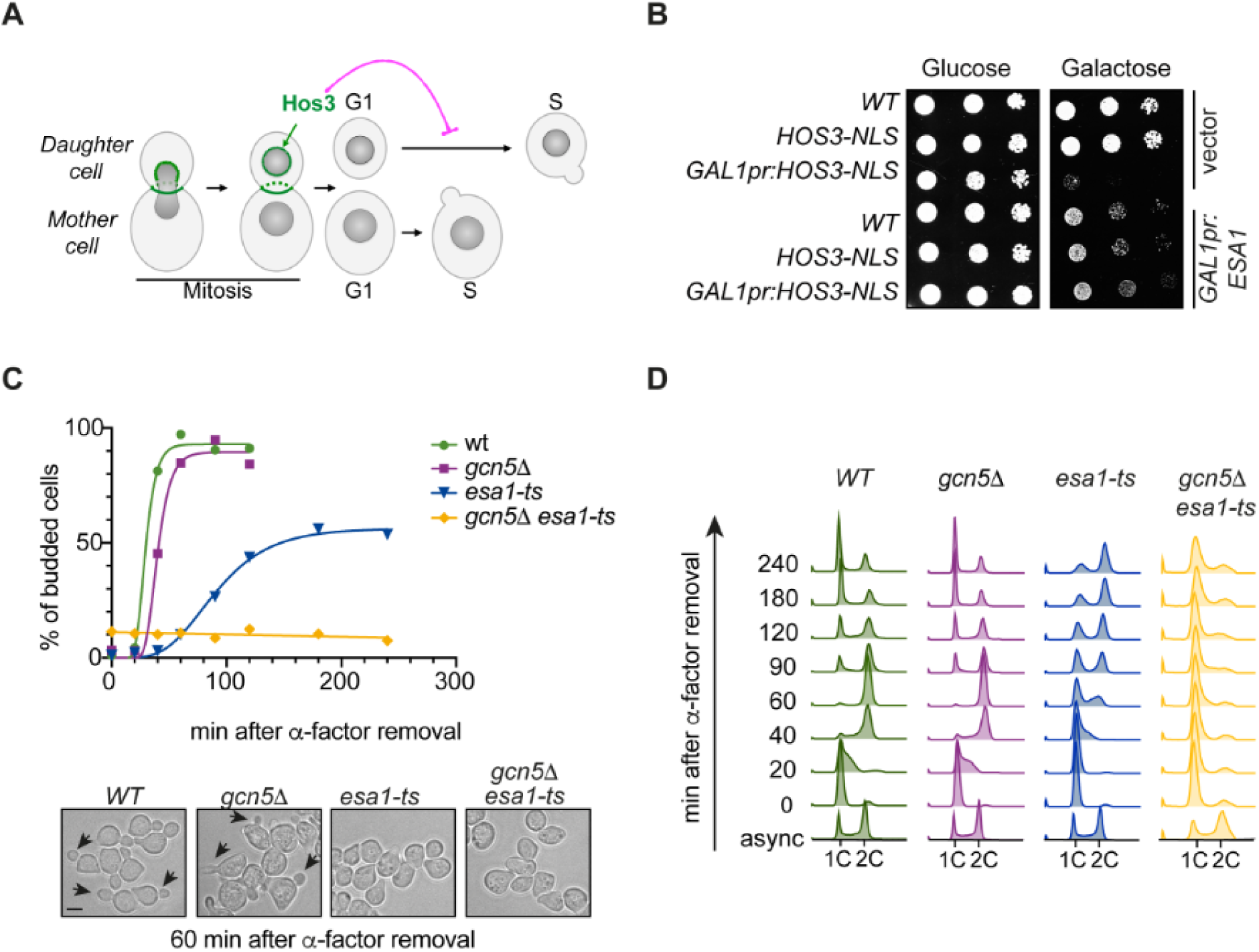
The KATs Esa1 and Gcn5 counteract the KDAC Hos3 to promote the G1/S transition. **(A)** Localization and function of the Hos3 deacetylase during mitotic division. Hos3 (in green) associates with the bud neck and with daughter-cell NPCs during nuclear migration into the bud. Hos3 delays the G1/S transition specifically in daughter cells through deacetylation of NPCs (Kumar et al., 2018). **(B)** Growth inhibition upon overexpression of *HOS3-NLS* is suppressed by overexpression of the KAT Esa1. 10-fold serial dilutions of the indicated strains transformed with an empty vector or the indicated plasmids, were spotted onto SC-Glu and SC-Gal medium and incubated at 25 °C for 3 days. Note that *HOS3-NLS* (under the control of the native *HOS3* promoter) does not affect cell growth. **(C)** *esa1-ts* and *gcn5Δ esa1-ts* mutants have bud emergence defects. (*Top*) Cells of the indicated strains were arrested in G1 by treatment with ɑ-factor for 2.5 h at 25 °C, shifted to 37 °C for 1 h and released from the G1 arrest at 37 °C. Cells were fixed at the indicated times and the presence of buds was assessed by microscopy. At least 200 cells were scored for each strain and time point. (*Bottom*) Bright field images of the indicated strains 60 min after the ɑ-factor washout. Arrowheads point to cell buds. Scale bar, 4 µm. **(D)** Inactivation of *ESA1* and *GCN5* delays DNA replication. Cells of the indicated genotypes were synchronised as in panel C, and DNA content was evaluated by flow cytometry. Numbers indicate time in minutes after the release. Experiments in C-D were repeated three times with similar results; one experiment is shown.

Here we show that the KAT Esa1 acetylates the nuclear basket component Nup60 to promote mRNA export and the G1/S transition. Furthermore, we demonstrate that Hos3-dependent deacetylation of Nup60 displaces mRNA export complexes from daughter-cell NPCs to inhibit Start. We propose that, in addition to modulating cell cycle entry and preventing premature division of daughter cells, this pathway regulates general mRNA export. Thus, the evolutionarily conserved NuA4 complex drives gene expression and cell cycle progression not only by acetylating chromatin and promoting transcription, but also by acetylation of the nucleoplasmic entrance to NPCs to facilitate export of nuclear mRNA, thereby dually controlling the gene expression state of the cell.

## Results

### The lysine acetyl-transferase Esa1 counteracts the Hos3 deacetylase and promotes cell cycle entry

To understand how NPC acetylation regulates the G1/S transition (Start), we sought to identify the lysine acetyl-transferases (KATs) counteracting the activity of the Hos3 deacetylase. Hos3 displays asymmetric distribution between mother and daughter cells in wild type *Saccharomyces cerevisiae*. Overexpression of a version of Hos3 fused to a nuclear localization signal (*GAL1pr-HOS3-NLS*) leads to targeting of Hos3 to mother and daughter cell nuclei, deacetylation of nucleoporins, and inhibition of cell proliferation (Kumar et al., 2018). We tested whether this inhibition could be relieved by overexpression of yeast KATs, including Eco1, Elp3, Esa1, Gcn5, Hat1, Hpa2, Hpa3, Rtt109, Sas2 and Spt10. Over-expression of Elp3, Gcn5 and Spt10 was toxic in wild-type cells, and therefore their potential role in opposing Hos3 could not be established using this assay (**Supplementary Figure S1A-B**). However, we found that of the remaining KATs, only Esa1 and Hat1 overexpression suppressed Hos3-NLS lethality (**Figure 1B** and **Supplementary Figure S1B**).

We next tested if inactivation of Esa1 or Hat1, alone or in combination, inhibits the G1/S transition, as would be expected of KATs counteracting the KDAC Hos3. We also tested the role of Gcn5, since its loss was previously reported to cause a mild delay in the G1/S transition (Kishkevich et al., 2019). The non-essential genes *HAT1* and *GCN5* were deleted, whereas Esa1 was inactivated using the well-characterised thermosensitive (ts) mutation *esa1-L254P* (hereafter called *esa1-ts*) (Clarke et al., 1999). Wild-type and mutant cells were arrested in G1 at 25 °C by addition of alpha factor, shifted to the restrictive temperature for *esa1-ts* (37 °C), and released from the cell cycle arrest by alpha factor removal. The fraction of S-phase cells at different times after alpha factor washout was determined by monitoring bud emergence. More than 95% of wild-type cells budded within 60 minutes of alpha factor removal, and *gcn5Δ* cells exhibited a 15-minute delay in budding as previously reported (Kishkevich et al., 2019). In contrast, budding was strongly delayed in *esa1-ts* cells: on average, only 25% of these cells had formed a bud after 60 minutes, and approximately 35% lacked a bud after 4 hours. Moreover, cells lacking both Esa1 and Gcn5 had stronger defects in budding than either single mutant: *esa1-ts gcn5Δ* cells remained unbudded after 4 hours of alpha factor washout (**Figure 1C**, **Supplementary Figures S2A and S2C**). Deletion of *HAT1* did not delay budding in either wild-type or *esa1-ts* backgrounds (**Supplementary Figure S2B**). Thus, *HAT1* does not play a role in Start and was not characterised further. DNA replication, assayed by flow cytometry, was also delayed in Esa1-deficient cells. Whereas most wild-type cells replicated their DNA 40 minutes after removal of alpha factor, replication was still incomplete after 4 hours in *esa1-ts* cells, and was undetectable in *esa1-ts gcn5Δ* (**Figure 1D**). In summary, Esa1 promotes budding and DNA replication, which are hallmarks of the G1/S transition. In the absence of Esa1, these functions can be partially compensated by Gcn5.

### Esa1 acts through Nup60 acetylation to promote Start

These data raised the possibility that Esa1, Gcn5 and Hos3 regulate the G1/S transition, at least in part, by modulating the acetylation level of shared target proteins. Proteomic studies have indicated that budding yeast nucleoporins are targeted by multiple KATs, including Esa1 and Gcn5, although the role of these modifications remained unclear (Henriksen et al., 2012; Downey et al., 2015). Furthermore, Hos3-dependent deacetylation of the nuclear basket component Nup60 lysine 467 (K467) is important for inhibition of Start in daughter cells (Kumar et al., 2018). Therefore, we investigated whether Esa1 and Gcn5 promote Start through Nup60 acetylation. Nup60-GFP was immunoprecipitated from cells expressing Esa1 or Gcn5 under the control of the inducible *GAL1* promoter, and its acetylation state was assayed with an anti-acetyl-lysine (AcK) antibody. This revealed increased Nup60 acetylation after addition of galactose in *GAL1pr-ESA1* and *GAL1pr-GCN5* cells (**Figure 2A**). We next tested if an acetyl-mimic form of Nup60 can restore the budding defect of Esa1-deficient cells. Lysine 467 of Nup60 was replaced with an asparagine (N) residue, whose biophysical properties of asparagine resemble those of acetylated lysine, to generate the acetyl-mimic Nup60-KN. Cells were released from a G1 block and their budding efficiency was determined as previously. Expression of Nup60-KN was not sufficient to restore budding in cells lacking both Esa1 and Gcn5 (**Supplementary Figure S3**), but it partially rescued the budding efficiency of the single mutant *esa1-ts* (**Figure 2B**). Thus, Esa1 promotes the G1/S transition in part by acetylation of Nup60 at lysine 467.

**Figure 2:**
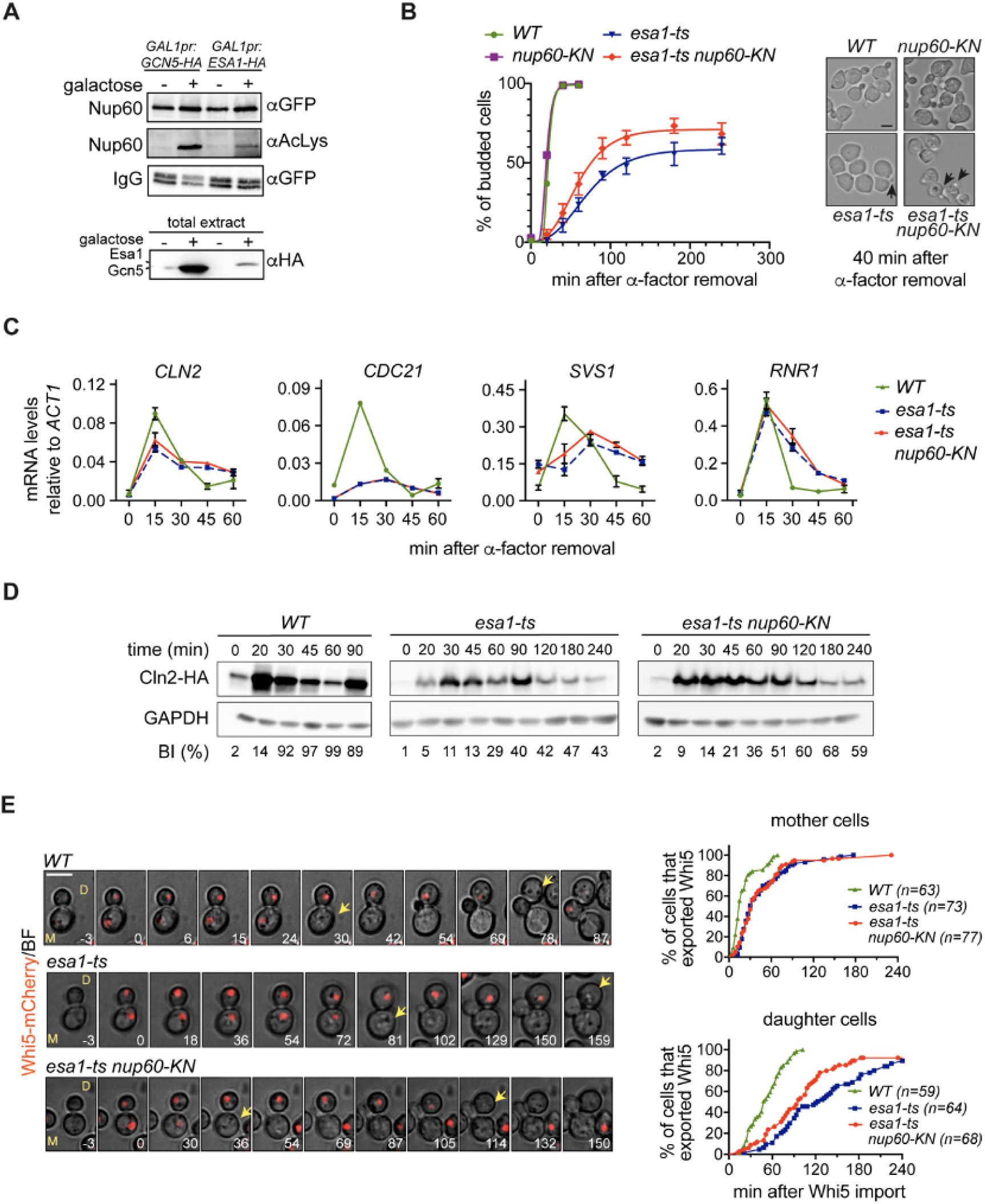
Acetyl-mimic Nup60 partially rescues the Start defects of *esa1-ts* cells. **(A)** Overexpression of Esa1 and Gcn5 KATs leads to increased acetylation levels of the nuclear basket nucleoporin Nup60. *(Top)* Nup60-GFP was immunoprecipitated from extracts of the indicated strains, and its acetylation state probed with anti-AcLys antibodies. *(Bottom)* Total extracts probed with anti-HA antibody to verify KAT overexpression. **(B)** *nup60-KN* partially rescues the budding defect of *esa1-ts* cells. *(Left)* Budding of cells of the indicated strains were determined as in Figure 1C. At least 200 cells were scored for each strain and time point. Data from 3 independent experiments is represented as mean and SEM. *(Right)* Bright field images of the indicated strains 40 min after the ɑ-factor washout. Arrowheads point to cell buds. Scale bar, 4 µm. **(C)** mRNA levels of *CLN2*, *SVS1*, *CDC21*, and *RNR1* were determined for cells of the indicated strains after G1 arrest and release at restrictive temperature, with samples collected at indicated times. Data from 3 independent experiments is represented as mean and SEM. **(D)** *nup60-KN* mutation partially rescues the delay in synthesis of the G1/S cyclin Cln2 in *esa1-ts* cells. Cells of the indicated strains were processed as in (B) and the amount of Cln2-HA protein at the indicated times was assessed by western blot. % of budded cells is indicated below for each corresponding strain and time point. Note the slow, inefficient budding in *esa1-ts* mutants. In *WT*, the reduction of Cln2 at 60 min and its increase at 90 min reflects the start of a second cycle, which is absent in *esa1-ts* cells. **(E)** *nup60-KN* partially rescues the Whi5 export defect of *esa1-ts* daughter cells. Composite of bright field and Whi5-mCherry (*left*) and quantification of Whi5 nuclear export (*right*) in mother (M) and daughter (D) cells of the indicated strains. Whi5 export (arrows) is delayed in *esa1-ts* mothers and daughters compared to *WT* (p < 000.1, Log-rank Mantel-Cox test); *esa1-ts nup60-KN* advances Whi5 export relative to *esa1-ts* in daughters (p=0.0105), but not in mothers (p > 0.05). 3 z-confocal slices spaced 0.5 µm were acquired every 3 min; maximum projections of selected timepoints are shown. Time is indicated in minutes; t=0 marks Whi5 nuclear import. Scale bar, 5 µm. *n* = number of cells, pooled from two independent experiments with similar results.

Start is marked by the transcription of hundreds of genes of the G1/S regulon, which are required for budding and DNA replication. Therefore, we next determined the mRNA levels of four representative regulon genes (*CLN2, CDC21*, *SVS1* and *RNR1*) in *esa1-ts* and *esa1-ts nup60-KN* synchronous cultures, using quantitative reverse transcription PCR (RT-qPCR). In agreement with the observed budding and DNA replication defects, transcription of G1/S genes was impaired in *esa1-ts* cells. In wild-type synchronous cultures, *CLN2, CDC21*, *SVS1* and *RNR1* are induced 15 minutes after removal of alpha factor, and their mRNA levels decrease as cells enter S phase (**Figure 2C**, *WT*). This pattern was disrupted in *esa1-ts* cells: *CLN2* was induced at lower levels and with slower kinetics than in wild type, whereas *CDC21* and *SVS1* mRNA levels did not oscillate during the experiment, and *RNR1* was induced at normal levels but its mRNA remained unusually stable (**Figure 2C**, *esa1-ts*). Together, these results indicate that Esa1 is required for coordinated induction of G1/S genes and bud emergence. However, the acetyl-mimic version of Nup60, which partially rescued budding efficiency of the *esa1-ts* mutant (**Figure 2B**), did not improve its transcriptional defects (**Figure 2C**, *esa1-ts nup60-KN*). To understand how Nup60-KN promotes budding of Esa1-deficient cells, western blotting was used to determine the protein levels of the G1/S cyclin Cln2 in *esa1-ts* and *esa1-ts nup60-KN*. Cln2 plays a critical role in driving activation of CDK in late G1 and robust, irreversible G1/S transition via positive feedback (Skotheim et al., 2008; Charvin et al., 2010). As expected, Cln2 protein synthesis occurred later and at lower levels in *esa1-ts* than in wild type cells released from a G1 block. Importantly, the delay in Cln2 protein synthesis was alleviated in *esa1-ts nup60-KN* (**Figure 2D** and **Supplementary Figure S4**). This suggests that Nup60 acetylation promotes Cln2 expression at the post-transcriptional level.

Finally, we examined the requirement for Esa1 and Nup60 acetylation in the G1/S transition of mother (M) and daughter (D) cells, using time-lapse microscopy of freely cycling cells. To determine the time of the G1/S transition in single cells, we monitored the nuclear localisation changes of the Whi5 transcriptional repressor, a G1 marker. Whi5 is imported into the nucleus in late anaphase and its export in G1, driven by CDK phosphorylation, marks the irreversible commitment to S phase (Costanzo et al., 2004; de Bruin et al., 2004; Charvin et al., 2010). In wild-type cells, nuclear export of Whi5-mCherry occurred first in mothers and later in daughters (median times, 15 min [M cells] and 59 min [D cells]; note that the duration of G1 phase in cells synchronised with alpha factor is not directly comparable with that of freely cycling cells) (**Figure 2E**, *WT* and **Supplementary Figure S5**). This dichotomy is due to both cell size control in small daughters and to size-independent mechanisms that delay Start specifically in daughters, including NPC deacetylation (Di Talia et al., 2007; Kumar et al., 2018). In *esa1-ts* cells, Whi5 export was markedly delayed in both M and D cells (median times: 30 min [M] and 123 min [D]; **Figure 2E**, *esa1-ts*). Furthemore, the presence of Nup60-KN partially restored the delay in Whi5 export caused by Esa1 inactivation, specifically in daughter cells (median times: 33 min [M] and 96 min [D]; **Figure 2E**, *esa1-ts nup60-KN*). Thus, Esa1 promotes Start in both mother and daughter cells, but constitutive Nup60 acetylation advances Start specifically in Esa1-deficient daughters. This is consistent with daughter-specific Nup60 deacetylation caused by asymmetric inheritance of the Hos3 KDAC (Kumar et al., 2018), and further supports the hypothesis that Esa1-dependent acetylation of Nup60 in mothers is reversed by Hos3 in daughter cells.

Together, these findings suggest two distinct roles for lysine acetylation in driving cell cycle commitment. First, a major function of Esa1 in promoting the G1/S transition is to drive the timely transcription of genes required for cell cycle entry. This is in keeping with an established role of Esa1 in promoting transcription of most yeast genes (Bruzzone et al., 2018). Second, Esa1 may play an additional positive role in Start, independently of transcription, that is mediated by Nup60 acetylation.

### Esa1 and Nup60 acetylation promote mRNA export

Nuclear basket components are required for efficient export of nuclear mRNA through their association with the Transcription and Export 2 (TREX-2) complex (Fischer et al., 2002). The finding that Nup60-KN promotes synthesis of Cln2 protein but not of its mRNA (**Figure 2C-D**) raises the possibility that Nup60 acetylation promotes mRNA export. We therefore tested whether the cell proliferation defect of *GAL1pr-HOS3-NLS* cells, in which Nup60 and other nucleoporins are deacetylated, can be alleviated by increased levels of mRNA export factors (**Figure 3A**). Indeed, we found that growth of *GAL1pr-HOS3-NLS* cells was restored by over-expression of the mRNA export receptors Mtr2 and Mex67, which escort mRNA molecules through the NPC (Strässer et al., 2000; Strawn et al., 2001) (**Figure 3B**). Likewise, overexpression of the scaffolding subunit of the TREX-2 complex, Sac3 (Fischer et al., 2002; Jani et al., 2014), also restored growth of *GAL1pr-HOS3-NLS* cells (**Figure 3C**). Note that a truncated Sac3 version was used, since overexpression of full-length Sac3 is toxic (**Supplementary Figure S6**). This further suggests that Hos3, possibly via Nup60 deacetylation, prevents cell proliferation by inhibiting export of nuclear mRNA.

**Figure 3:**
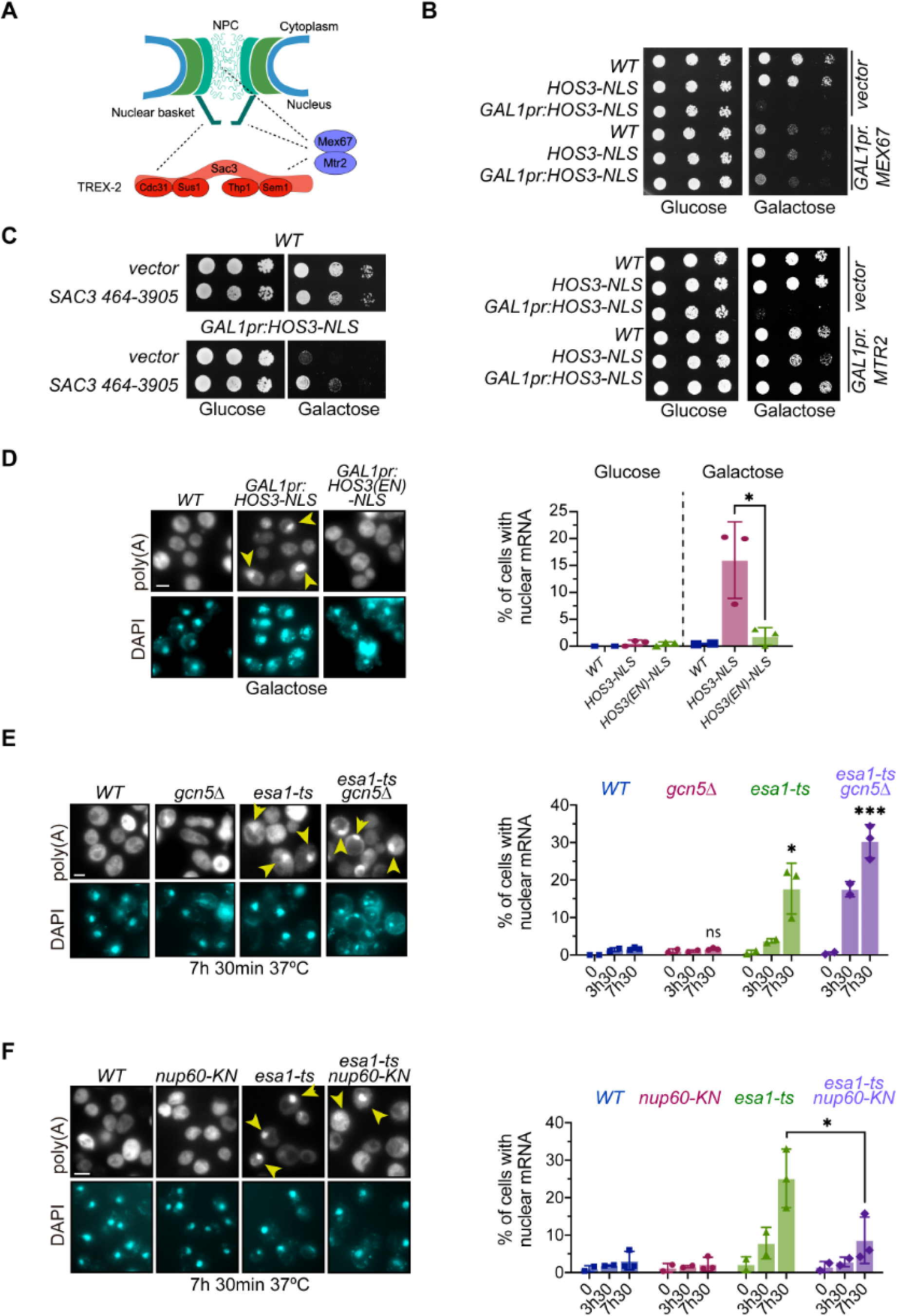
Nuclear export of mRNA is inhibited by the KDAC Hos3, and promoted by the KATs Esa1, Gcn5, and by Nup60 acetylation. **(A)** Illustration of the physical interactions (dashed lines) of the NPC with mRNA export factors. **(B)** Overexpression of *MEX67* and *MTR2* rescues the lethality of Hos3-NLS overexpression. 10-fold serial dilutions of the indicated strains transformed with the indicated plasmids spotted onto SC-Glu and SC-Gal medium and incubated at 25 °C for 3 days. **(C)** A high-copy plasmid containing *SAC3* rescues the lethality of Hos3-NLS overexpression. Strains carrying a high copy plasmid containing the *SAC3* ORF (nucleotides 467-3905), or the empty vector, were grown as in (B). **(D)** Overexpression of Hos3-NLS promotes nuclear accumulation of mRNA. (*Left*) Cultures of the indicated strains were treated with galactose overnight to induce *HOS3-NLS* expression, cells were fixed, and FISH was performed using Cy3-Oligo(dT). Arrows point to polyadenylated RNA in the nucleus, which was visualized by DAPI staining. (*Right*) The fraction of cells with nuclear mRNA accumulation was determined for the indicated strains and conditions. **(E)** Inactivation of Esa1 impairs export of poly(A) RNA. (*Left*) Cultures of the indicated strains were incubated at 37 °C. (*Right*) The fraction of cells with nuclear mRNA accumulation, marked with arrows, was determined for the indicated strains and conditions as in (D). **(F)** *nup60-KN* mutation partially rescues the mRNA export defects of *esa1-ts.* Cells of the indicated strains were processed as in (E). In (D-F), graphs show the mean and s.d. from three biological replicates.*, p ≤ 0.05; ***, p ≤ 0.001; ns, p > 0.05, two-tailed unpaired t-test. At least 200 cells were scored for each time-point and condition. Scale bar, 4 µm.

To directly test whether Nup60 deacetylation inhibits mRNA export, we imaged polyadenylated mRNA by fluorescence in situ hybridization (FISH) using a poly-dT probe. This was done in wild type and in cells in which Nup60 acetylation was reduced by over-expression of nuclear Hos3 (*GAL1pr:HOS3-NLS*), or by inactivation of Esa1 and Gcn5 (*esa1-ts*, *gcn5Δ*). As expected, all wild-type cells showed mRNA localization diffusely in both nucleus and cytoplasm. In contrast, approximately 15% of cells exhibited nuclear mRNA accumulation upon induction of *GAL1pr:HOS3-NLS* (**Figure 3D** and **Supplementary Figure S7A)**. Nuclear accumulation of mRNA was dependent on Hos3 KDAC activity, as it was not observed upon overexpression of a catalytically inactive mutant (Hos3^EN^-NLS) (**Figure 3D**). Furthermore, inactivation of Esa1 (*esa1-ts*) led to accumulation of nuclear mRNA in up to 20% of cells, and this fraction rose to 30% in the double mutant *esa1-ts gcn5Δ* (**Figure 3E** and **Supplementary Figure S7B**). Notably, deletion of *GCN5* alone did not cause nuclear mRNA accumulation, suggesting this KAT does not play an important role in promoting mRNA export. The Esa1 function in RNA export appeared to be specific for mRNA, since depletion of Esa1 (alone or in combination with Gcn5) did not affect export of ribosomal RNA (**Supplementary Figure S8).** Importantly, the fraction of Esa1-deficient cells with nuclear mRNA accumulation was significantly reduced in *esa1-ts* cells carrying the acetyl-mimic Nup60 mutation (**Figure 3F**). These results suggest that acetylation and deacetylation of Nup60, mediated by Esa1 and Hos3 respectively, regulate mRNA export.

### Nup60 acetylation recruits the TREX-2 complex to the nuclear basket to promote Start

We previously found that constitutive nuclear localisation of Hos3 (Hos3-NLS) reduces the amount of NPC-associated mRNA export factors such as Sac3, that localise to the nuclear periphery (Kumar et al., 2018). This suggests that in wild type cells, Nup60 acetylation facilitates the recruitment of Sac3 to NPCs and this is inhibited in G1 daughter cells via Hos3-dependent Nup60 deacetylation. To test this prediction, we measured the nuclear intensity of GFP-tagged Sac3 relative to that of the structural NPC component Nup49-mCherry in mother (M) and daughter (D) nuclei immediately after cytokinesis. Interestingly, loss of Hos3 (*hos3Δ*) lead to an increase of Sac3 nuclear localisation in D cells, and expression of acetyl-mimic Nup60 (*nup60-KN*) caused an increase of nuclear Sac3 in both M and D cells (**Figure 4A**, nucleus). Importantly, total Sac3 intensity was not affected by mutations in *HOS3* or *NUP60* (**Figure 4A**, whole cell). We conclude that Hos3 and Nup60 deacetylation reduce the enrichment of Sac3 in G1 daughter cell nuclei. We then tested whether Esa1 promotes the localisation of Sac3 to the nuclear basket. To avoid potential confounding effects due to Esa1 inactivation during S phase, cells were treated with the microtubule polymerisation inhibitor nocodazole at 25 °C, to arrest *esa1-ts* cells in mitosis in the presence of Esa1 function. Nocodazole was then removed, cells were shifted to 37 °C to inactivate Esa1, and the nuclear intensity of Sac3-GFP, normalised to that of Nup49-mCherry, was determined in the following G1. This revealed that nuclear enrichment of Sac3 is significantly reduced in both M and D *esa1-ts* cells, while total cellular levels of Sac3 were not affected (**Figure 4B**). Together, these data indicate that Nup60 acetylation, driven by Esa1 and inhibited by Hos3 in G1 daughter cells, is important for perinuclear recruitment of the mRNA export factor Sac3.

**Figure 4:**
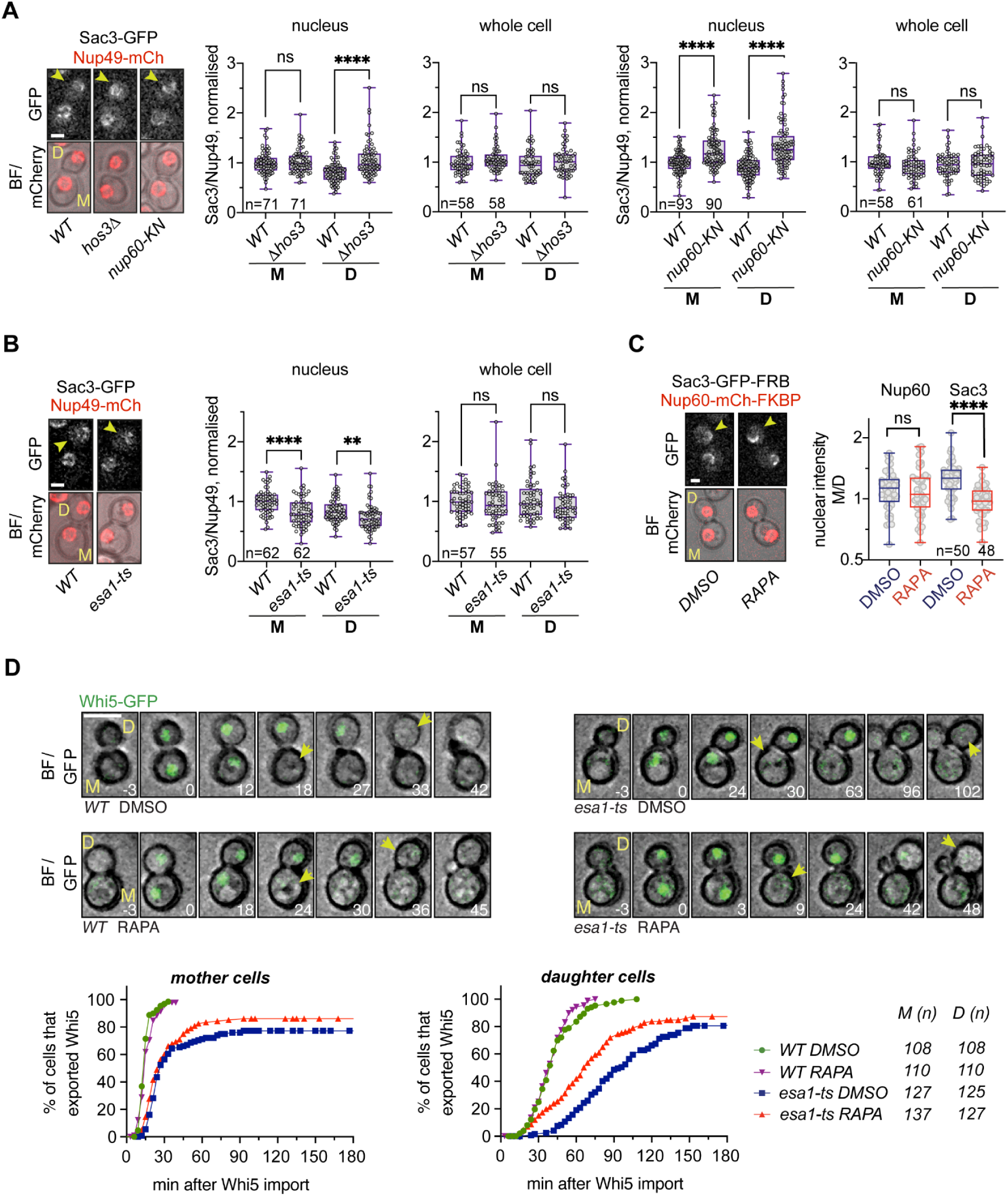
Nup60 deacetylation in daughter cells displaces Sac3 from NPCs and delays Start. **(A)** Depletion of Hos3 or expression of acetyl-mimic Nup60 (*nup60-KN*) increases the nuclear localisation of Sac3. Cells of the indicated strains were imaged by time-lapse microscopy and the fluorescence levels of the indicated proteins were determined after cytokinesis. The NPC component Nup49 was used as a control for Nuclear Pore Complex protein levels. Fluorescence intensity was measured in sum projections of whole-cell Z-stacks, by segmentation of either the nuclear area in the mCherry channel, or of the whole cell in the brightfield channel. The ratio of Sac3/Nup49 intensities was then normalised relative to the median intensity of wild-type mothers. **(B)** Inactivation of Esa1 decreases Sac3 nuclear levels. Wild-type (*WT*) and *esa1-ts* cells were arrested in mitosis by treatment with nocodazole at 25 °C, shifted to 37 °C, released from the mitosis block in fresh medium at 37 °C, and imaged by time-lapse microscopy. Fluorescence levels were quantified in G1 as in (A). In (A-B), arrowheads point to daughter cells. Scale bar, 2 µm. **(C)** Rapamycin-dependent dimerisation abolishes Sac3 mother/daughter asymmetries. *NUP60-mCherry-FKBP SAC3-GFP-FRB* cells were incubated with rapamycin (RAPA) to trigger FRB-FKBP heterodimerization, or with DMSO as control. Fluorescence levels were quantified in G1 cells as in (A), 15 to 30 minutes after addition of the drug. In (A-C), boxes include 50% of data points, the line represents the median and whiskers extend to maximum and minimum values. Individual cells are shown in grey. ****, p ≤ 0.0001; ns, p > 0.05, two-tailed unpaired t-test. **(D)** Sac3 anchoring to the nuclear basket advances Start in *esa1-ts* daughter cells. Composite of bright field and Whi5-mGFP (*top*) and quantification of Whi5 nuclear export timing (*bottom*) in wild-type (*WT*) and *esa1-ts* mother (M) and daughter (D) cells treated with either rapamycin (RAPA) or DMSO and expressing Nup60-FRB and Sac3-mCherry-FKBP. Sac3 anchoring to Nup60 does not alter Whi5 export timing in *WT* mother or daughter cells (DMSO vs RAPA, p > 0.05, Log-rank Mantel-Cox test), but it advances Whi5 export in *esa1-ts* daughters (p = 0.0001). Whi5 export efficiency was slightly improved also in mother cells (p = 0.0374). 8 z-confocal slices spaced 0.4 µm were acquired every 3 min; maximum projections are shown. Time is indicated in minutes; t=0 marks Whi5 nuclear import. Scale bar, 5 µm. *n* = number of cells, pooled from three independent experiments with similar results.

Sac3 is tethered to the nuclear pore basket, and this localisation is required for its mRNA export function (Fischer et al., 2002). To test if Esa1 promotes Start by targeting Sac3 to NPCs, we asked whether artificially anchoring Sac3 to the nuclear basket is sufficient to rescue their Start delay in Esa1-deficient cells. We used the anchor-away system, which triggers dimerisation of FK506-binding protein (FKBP) and FKBP-rapamycin-binding (FRB) in the presence of rapamycin (Gallego et al., 2013) to anchor Sac3-FKBP and Nup60-FRB protein fusions. Consistent with results in Figure 4A, the distribution of fluorescently labeled Sac3-FRB and Nup60-FKBP was biased towards mother cells, with a higher bias for Sac3 than for Nup60 (p<0.005, unpaired t-test) (**Figure 4C**). Addition of rapamycin did not alter the accumulation of Nup60-FKBP in mother cells. In contrast, rapamycin altered the asymmetric localisation of Sac3-FRB: whereas Sac3 accumulated preferentially in mother cell nuclei in DMSO-treated cells, it was partitioned equally to M and D nuclei in the presence of rapamycin (**Figure 4C**). Thus, rapamycin increases the localisation of Sac3-FRB in daughter cells. G1 duration was then determined by time-lapse microscopy of *esa1-ts* cells expressing Whi5-GFP, as in Figure 2E, in the presence of either rapamycin or DMSO. Rapamycin slightly increased Whi5 export efficiency in *esa1-ts NUP60-FRB SAC3-FKBP* mother cells, and specifically advanced Whi5 export in daughter cells (**Figure 4D**). This is consistent with Hos3-dependent Nup60 deacetylation in daughter cells (Kumar et al., 2018). We conclude that Esa1 and Nup60 acetylation promote Start, at least in part, by targeting Sac3 to the nuclear basket, where it mediates mRNA export. Consistent with the requirement of mRNA export to trigger Start, we find that inactivation of the essential mRNA export factor Mex67 is sufficient to prevent entry into S phase (**Supplementary Figure S9**).

### Nup60 acetylation regulates expression of the inducible *GAL1* gene

Our results indicate that Nup60 deacetylation inhibits mRNA export (**Figure 3**) and reduces the NPC recruitment of Sac3 to delay Start in daughter cells, presumably by inhibiting the export of mRNAs required for S phase (**Figure 4**). Next, we asked whether Nup60 acetylation can also affect the expression of genes that are not required for the G1/S transition, such as the inducible galactokinase (*GAL1*) gene. To measure *GAL1* expression, fast-folding GFP (*sfGFP*) was placed under the control of the *GAL1-10* promoter (*GAL1pr*) and inserted next to the endogenous *GAL1* locus. Thus, measuring GFP fluorescence with time lapse-confocal microscopy allows tracking of *GAL1* expression in single cells. Wild type and Nup60 acetyl-mimic (*nup60-KN*) were placed in a microscope chamber, and *GAL1* expression was induced with galactose (**Figure 5A**). GFP fluorescence appeared earlier and increased to higher levels in *nup60-KN* than in wild-type cells, suggesting that Nup60 acetylation promotes *GAL1* expression (**Figure 5B** and **Supplementary Figure S10**). Notably, Nup60 levels were equivalent throughout all experiments, suggesting that Nup60 acetylation is unlikely to affect gene expression through changes in Nup60 stability (**Supplementary Figure S11**). Because Nup60 acetylation is inhibited by Hos3 in daughter cells, we tested whether *GAL1* expression occurs with different strength in mothers and daughters. Indeed, *GAL1* expression levels were higher in wild-type mother cells than in their daughters, and these differences were absent in cells lacking Hos3 or expressing Nup60-KN (**Figure 5C**). We conclude that acetylation of Nup60 in mother cells promotes *GAL1* expression, whereas its deacetylation in daughter cells inhibits expression. Expression of *GAL1* was slightly increased in the double mutant *hos3Δ nup60-KN* relative to either *hos3Δ* and *nup60-KN* single mutants (**Figure 5B-C**). This indicates that Hos3 inhibits *GAL1* expression in daughters largely, but not entirely through deacetylation of Nup60 at Lys 467. Furthermore, forcing the symmetric distribution of the mRNA export factor Sac3 to the nuclear basket of mother and daughter nuclei using the FRB-FKBP system (as in **Figure 4C**) increased *GAL1* expression specifically in daughter cells (**Figure 5D-E**). Thus, Hos3-dependent deacetylation of Nup60 inhibits *GAL1* expression through Sac3, which regulates the export and/or synthesis of *GAL1* mRNA.

**Figure 5:**
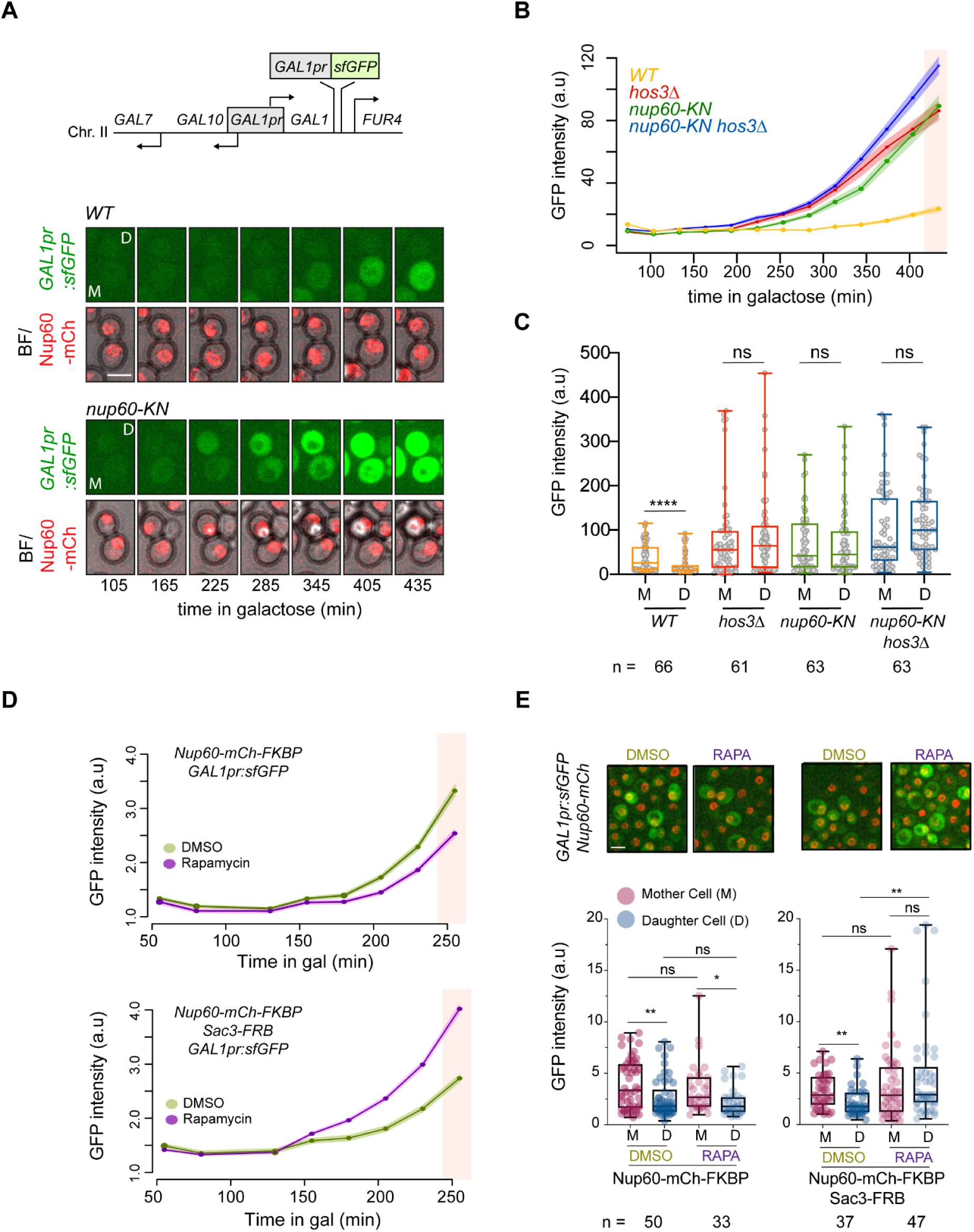
Daughter-cell specific Nup60 deacetylation inhibits *GAL1* expression through displacement of Sac3 from NPCs. **(A)** (*Top*) The *GAL1pr:sfGFP* reporter was integrated on Chr. II between the *GAL1* and *FUR4* loci. (*Bottom*) Time Lapse microscopy of *WT* and *nup60-KN* cells expressing *GAL1pr:sfGFP* and Nup60-mCherry at the indicated times of galactose induction. Scale bar, 4 µm. **(B)** Depletion of Hos3, and expression of acetyl-mimic Nup60 (*nup60-KN*) enhance *GAL1* expression. WT, *hos3Δ, nup60-KN* and *nup60-KN hos3Δ* cells were shifted to galactose and imaged by time lapse microscopy to monitor *GAL1pr:sfGFP* expression during 7 hours. Nuclear fluorescence was scored by segmentation of the nuclear area in the mCherry channel and mean fluorescence of nuclear GFP and Nup60-mCherry was quantified from sum projections of whole-cell Z-stacks at the indicated times. At least 200 cells were scored for each strain and time point. Shaded areas indicate the SEM. **(C)** GFP intensity of mother/daughter pairs were quantified as in B at 425 min after galactose addition (pink shaded area in B). **(D)** Sac3 anchoring to Nup60 advances *GAL1* expression. Cells of the indicated strains were incubated with galactose in the presence of either rapamycin (RAPA) to induce FRB-FKBP heterodimerization, or DMSO as control. *GAL1pr:sfGFP* expression was monitored over time as in B. **(E)** (*Top*) Representative images of the indicated cells in rapamycin or DMSO, 250 min after galactose addition. Scale bar 4 µm. (*Bottom*) Mother/daughter pairs were quantified as in B at 250 min after galactose addition (pink shaded area in D). In (C and E), boxes include 50% of data points, the line represents the median and whiskers extend to maximum and minimum values. ***, p ≤ 0.001; **, p ≤ 0.01; *, p ≤ 0.05; ns, p > 0.05, two-tailed paired t-test for M-D comparisons, unpaired for comparisons between strains..

## Discussion

Our data indicate that in budding yeast, the lysine acetyltransferase subunits of the NuA4 complex (Esa1) and to a lesser extent, the SAGA complex (Gcn5), are required for cell cycle entry. Given that the G1/S transition depends on the coordinated expression of hundreds of genes, the requirement of NuA4 and SAGA components in this cell cycle transition is in line with their common role as transcriptional coactivators with known functional overlaps. Indeed, NuA4 and SAGA complexes share the targeting subunit Tra1 (Helmlinger and Tora, 2017) and are thought to drive gene expression mainly through acetylation of histone H3 (for SAGa) and H4 (for NuA4) and subsequent chromatin decompaction (Sterner and Berger, 2000; Lee and Workman, 2007). However, we demonstrate that Esa1 (but not Gcn5) is required not only for gene transcription but also for mRNA export. Importantly, expression of an acetyl-mimic Esa1 substrate, the nuclear pore component Nup60, partially alleviates the G1/S transition and mRNA export defects of Esa1-defective cells. Thus, we propose that Gcn5 and Esa1 have overlapping roles in transcription of G1/S genes, and that Esa1 is also involved in promoting mRNA export by acetylation of Nup60. To our knowledge, this is the first time that a KAT is linked to post-transcriptional regulation of gene expression by modification of NPCs. Esa1 may acetylate other NPC components in addition to Nup60 (Henriksen et al., 2012; Kumar et al., 2018). Importantly, we show that Esa1 activity and Nup60 acetylation facilitate the nuclear enrichment of the TREX-2 complex scaffolding subunit Sac3, which promotes mRNA export at the nucleoplasmic side of the nuclear pores, thereby driving mRNA export and cell cycle entry (**Figure 6**). Our finding that Nup60 acetylation promotes the Sac3-dependent expression of *GAL1*, which is not required for cell cycle entry, further suggests that Esa1 and Nup60 acetylation are part of a pathway promoting general mRNA export during the entire cell cycle.

**Figure 6:**
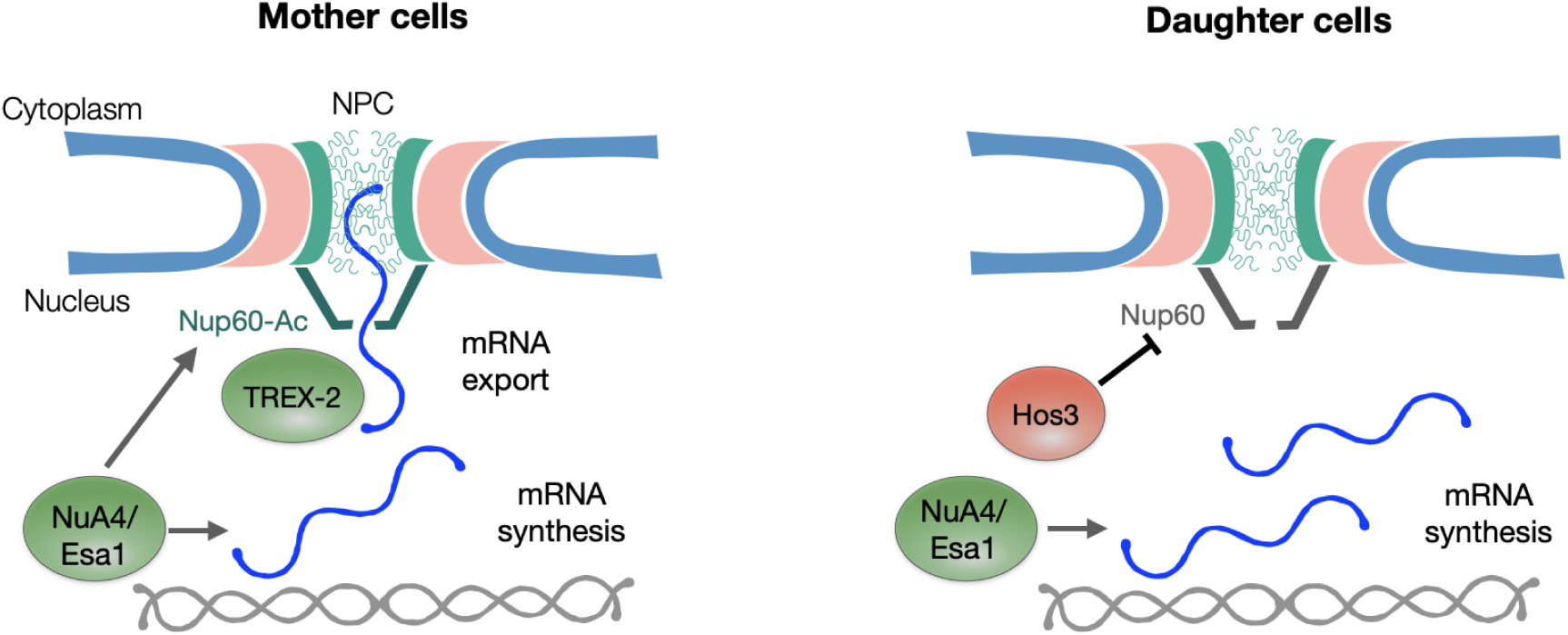
Esa1 coordinates mRNA synthesis and export during the G1/S transition though Nup60 acetylation. In mother cells, Esa1 promotes both mRNA synthesis and export. Mechanistically, mRNA export is promoted by acetylation of Nup60, which increases the association of the mRNA export factor TREX-2 to the nuclear pore basket. In daughter cells, Hos3 deacetylates Nup60, which reduces TREX-2 association with the NPC, and thus mRNA export. Inhibition of Nup60 acetylation in daughter cells contributes to their longer G1 phase, possibly by delaying the export of mRNAs required for entry into S phase, and inhibits the expression of the *GAL1* gene in response to galactose.

Our results also indicate that Esa1-dependent mRNA export, which may be constitutively active in mother cells, is specifically inhibited in daughter cells to restrain their G1/S transition. This inhibition is caused by the KDAC Hos3, which deacetylates Nup60 in G1 daughters, contributing to their prolonged G1 phase and enforcing cell size control. Nup60 acetylation in daughter cells is restored in S phase, presumably due to removal of Hos3 from NPCs in G1 (Kumar et al., 2018), thus allowing resumption of normal mRNA export in daughter cells later in the cell cycle. These observations reveal an additional level of control of the G1/S transition, which is also regulated by the differential scaling of Start inhibitors and activators with cell size (Schmoller et al., 2015; Chen et al., 2020) and by the daughter-specific inheritance of transcriptional regulators (Di Talia et al., 2009; Kumar et al., 2018). Notably, our data suggest that gene expression at the G1/S transition is controlled not only through transcription, but also at the level of mRNA export. We speculate that the Hos3-Nup60 pathway downregulates mRNA export during G1, because Nup60 deacetylation is largely restricted to this cell cycle phase. Moreover, it is possible that Nup60 deacetylation does not specifically inhibit expression of the G1/S regulon, because Hos3 also inhibits expression of the nutrient-responsive *GAL1* gene.

Esa1-deficient yeast cells arrest in late S-phase or early mitosis in a manner dependent on the DNA damage checkpoint (Clarke et al., 1999). This cell cycle arrest probably masked the role of Esa1 in earlier cell cycle stages, which we reveal here through analysis of both synchronised populations and freely cycling cells. Moreover, our results raise the possibility that DNA damage in *esa1-ts* cells may stem (at least in part) from replicative stress during S-phase caused by inefficient synthesis and/or export of G1/S mRNAs. Supporting this hypothesis, DNA replication (although delayed) seems to proceed in many unbudded *esa1-ts* cells (Figure 1C-D). In addition, lack of Esa1 activity is associated with aberrant nucleolar fragmentation (Clarke et al., 1999). Although the molecular basis of the nucleolar defect in Esa1-defective cells is unclear, it is interesting that nucleolar fragmentation is also observed after inactivation of mRNA export factors, provided that mRNA synthesis is ongoing (Kadowaki et al., 1994; Schneiter et al., 1995). Thus, nucleolar fragmentation after inactivation of Esa1 may be caused by abnormal accumulation of mRNA in the cell nucleus.

mRNA export factors, including Mex67 and Sac3, contribute to NPC tethering of active yeast genes, and may contribute to their optimal expression (Cabal et al., 2006; Dieppois et al., 2006; Brickner et al., 2019). Our data indicate that Esa1 promotes the perinuclear enrichment of Sac3 but whether this localisation affects the interaction of chromosomal loci with NPCs is not known. Interestingly, the *CLN2* locus interacts with NPCs specifically in G1 and this interaction is stabilised by Nup60 deacetylation in daughter cells (Kumar et al., 2018). It will be of interest to establish how NPC tethering of G1/S genes such as *CLN2* is related to the elongation and export state of their respective mRNAs, and whether stabilisation of gene tethering by Nup60 deacetylation corresponds to primed or active transcriptional conditions.

Whether the role of NuA4 in NPC acetylation and mRNA export is evolutionarily conserved is not known. As in yeast, the TREX-2 complex is associated with the nuclear pore basket in human cells (Umlauf et al., 2013) and promotes export of a subset of mRNAs together with the basket component TPR (Wickramasinghe et al., 2010, 2014; Aksenova et al., 2020). Furthermore, human nucleoporins are acetylated, including TPR and the Nup60 homologue Nup153 (Choudhary et al., 2009), and TPR physically interacts with Tip60/KAT5, the mammalian homologue of Esa1 (Chen et al., 2013); however the physiological relevance of these modifications and interactions has not been determined. Our findings raise the possibility that mammalian nucleoporins represent a novel category of substrates for KATs and for the multiprotein complexes in which these enzymes reside, with important roles in gene expression. Identification of the molecular mechanisms by which KATs such as NuA4 and SAGA regulate mRNA export in mammalian cells, by acetylation of non-histone proteins such as nucleoporins, remains therefore a key subject for future studies.

In summary, these data reveal a novel role in mRNA export for the evolutionarily conserved KAT-containing coactivator complex, NuA4. They also demonstrate that differences in Nup60 acetylation determined by the interplay between a KAT (in mother cells) and a KDAC (in daughter cells) allow the modulation of mRNA export capabilities of NPCs in different cell types, shaping their gene expression and cell proliferation profiles. Furthermore, our findings on the regulation of *GAL1* expression by Nup60 acetylation indicate that differences in NPC acetylation between mother and daughter cells contribute to the development of heterogeneous gene expression responses among a population of genetically identical cells. This type of phenotypic variability (often interpreted as a bet-hedging strategy) could provide a growth advantage for a clonal population upon sudden changes in environmental conditions (Veening et al., 2008). Thus, while specifically assessing the role of NPC acetylation in cell cycle entry, our study raises the possibility that acetylation of nuclear pores and regulation of mRNA export define an important regulatory step in cell identity establishment. Analogous mechanisms may also contribute to cell differentiation during the development of multicellular organisms.

## Acknowledgements

We are grateful to the IGBMC Imaging Centre and Flow Cytometry Facility, Kenny Schumacher for assistance with RT-qPCR; Didier Devys, Gabriel Neurohr, Snezhana Oliferenko, Laszlo Tora and all members of the Mendoza lab for helpful comments; and to Life Science Editors for editorial assistance. This study was supported by Ministerio de Ciencia e Innovación, Spain, and co-financed by the European Regional Development Fund from the European Union, grant numbers BFU2017-88692 and PID2020-119793GB-I00 to JCI, by the Fondation ARC pour la recherche sur le cancer www.fondation-arc.org and “Équipe FRM” EQU202003010561 to MM, and by the grant ANR-10-LABX-0030-INRT, which is a French State fund managed by the Agence Nationale de la Recherche under the frame programme Investissements d’Avenir ANR-10-IDEX-0002-02 to the IGBMC. MGA was a Postdoctoral Fellow from the Generalitat Valenciana (APOSTD/2017/094) and is a recipient of a Juan de la Cierva Incorporación Fellowship (IJC2018-036206-I) from the Ministerio de Ciencia e Innovación, Spain.

## Materials and Methods

### Strains, plasmids and cell growth

*Saccharomyces cerevisiae* strains are derivatives of S288c or BY4741 except when indicated (**Supplementary Table S1**). Gene deletions and insertions of C-terminal tags were generated by standard one-step PCR-based methods (Longtine et al., 1998; Janke et al., 2004). The acetyl-mimic *nup60-KN* mutant was generated using CRISPR/Cas9 to replace the acetylated lysine 467 by asparagine, as described (Kumar et al., 2018). The *esa1-ts* thermosensitive strain carrying the *L254P* mutation (Clarke et al., 1999) linked to the kanamycin-resistance cassette *KanMX* (Li et al., 2011) was used to obtain the *esa1-ts-kanMX* cassette and integrate the *ts* allele in the corresponding S288c-derived strains. The *mex67-ts* strain is a gift from Dr. M. del Olmo (Estruch et al., 2009).

pBG1805 2µ multicopy plasmids expressing KATs, Mtr2, Mex67 or Sac3 under the control of the *GAL1* promoter and carrying a HA C-terminal tag (Gelperin et al., 2005) were obtained from the Yeast ORF collection (Thermo Scientific Open Biosystems, YSC3868). pGP565 2µ multicopy plasmids (Jones et al., 2008) containing Sac3, Sus1, Cdc31, Sem1 and Thp1 are a gift from Dr. S. Leon (Yeast Genomic Tiling Collection, Open Biosystems). The pUC57 vector containing the superfolder (sf) GFP fused to the CLN2-PEST degron under the control of *GAL1pr* (AP2) is described in (Goulev et al., 2019) and the Rpl25-GFP plasmid is a gift from Dr. H. Schmidt.

Cells were grown in exponential conditions (below OD_600_=1) at 25 °C in standard yeast extract-peptone-dextrose medium supplemented with adenine 70 μg/ml (YPDA) or synthetic complete (SC) medium with 2% glucose, 2% raffinose (SC-Raf) or 2% galactose (SC-Gal). Where indicated, cells were incubated in the presence of 15 μg/ml nocodazole, 15 μg/ml α-factor, or transferred to 37 °C. For growth assays in solid media, 10-fold serial dilutions of exponential cultures were spotted onto SC-Glu and SC-Gal medium and incubated at 25 °C for 3 days.

For G1 arrest, exponential cells growing in YPDA medium were synchronized with 15 μg/ml α-factor (GenScript Cat. No:RP01002) for 2h at 25 °C, supplemented with additional α-factor (5 μg/ml) and incubated 30 min more at 25 °C. Then cells were shifted to 37 °C during 1 h, washed 3 times with pre-warmed YPDA and released in fresh pre-warmed YPDA medium at 37 °C. For the nocodazole arrest, cells were incubated during 2h with nocodazole 15 μg/mL, washed 3 times and then released in fresh pre-warmed YPDA medium at 37 °C.

For the analyses of *GAL1pr-*driven expression of the sfGFP reporter, the *GAL1pr:sfGFP-CLN2-PEST* cassette was integrated between the *GAL1* and *FUR4* loci using oligonucleotides GALsfGFP-Fw (5’-AAAGTCATTTGCGAAGTTCTTGGCAAGTTGCCAACTGACGtACGGATTAGAAGCCGCC GA-3’) and GALsfGFP-Rv (5’-AGGACAAAAAGTTTCAAGACGGCAATCTCTTTTTACTGCAATGGttagaaaaactcatcgag-3’) and the AP2 plasmid as a PCR template. For induction of the *GAL1pr:sfGFP* reporter (Figure 5), exponential cells were grown in glucose, washed 3 times with SC-Gal and resuspended in SC-Gal (2% galactose, 0.1% glucose) for time-lapse imaging. For Nup60-GFP IP assays (Figure 2A), cells were grown in glucose until exponential phase, diluted and incubated in SC-Raf (2% raffinose, 0.1% glucose) overnight until exponential phase and 2% galactose was added to induce *GAL1pr:GCN5-HA* and *GAL1pr:ESA1-HA* expression for 2 h.

For tethering of Sac3 to Nup60, we used inducible dimerization of FK506-binding protein (FKBP) and FKBP-rapamycin binding (FRB) domain, as described in (Gallego et al., 2013). Sac3 and Nup60 were tagged at the C-terminus. The background of the anchoring strains harbors the *tor1-1* mutation and lacks the endogenous FPR1 gene rendering growth insensitive to rapamycin. Exponential cells were incubated with 20 μM rapamycin (Sigma-Aldrich Cat. No R8781) throughout the imaging experiment. Association of Sac3-GFP-FRB with Nup60-mCherry-FKBP was checked by measuring the disruption of Sac3 asymmetries within 15-30 min of rapamycin addition. Similar results were obtained for Sac3-mCherry-FKBP and Nup60-FRB (Figure 4D). For the induction of *GAL1pr:sfGFP*, cells were grown in glucose, washed 3 times with SC-Gal and shifted to SC-Gal (2% galactose, 0.1% glucose) with rapamycin or DMSO for the time-lapse imaging (Figure 5D-E). For inactivation of *esa1-ts*, cells were imaged at 37 °C at the moment of rapamycin or DMSO addition (Figure 4D).

### Fluorescence microscopy

For time-lapse microscopy, cells were grown overnight in 50 ml flasks containing 10 ml of SC medium at 25 °C, then diluted to OD600 = 0.1-0.3 in fresh medium, grown at least for 4 hours to mid-log phase and plated in minimal synthetic medium on concanavalin A–coated (Sigma-Aldrich) Lab-Tek chambers (Thermo Fisher Scientific). Time-lapse imaging was performed using a confocal spinning disk (Nikon, Garden City, NY) equipped with Yokogawa CSU-X1 confocal scanner, temperature-control Tokai Hit Stage Top Incubator, a z-stepper and an automated stage, controlled by MetaMorph software. Images were acquired with an HCX PL APO 100X objective and a Photometrics Prime 95B camera. Time-lapse series of 4 μm stacks spaced 0.2-0.3 μm were acquired every 2-4 min. For Sac3-GFP a maximum of 13 z-stacks were taken, for Whi5-GFP 8 z-stacks spaced by 0.4 μm, and in case of Whi5-mCherry due to low protein abundance and poor fluorophore stability 3 z-stacks spaced by 0.5 μm were used. Red channel images (Whi5-mCherry) were subject to Gaussian blur to remove noise (radius = 2 px). The Images were processed and analyzed on 2D maximum or sum projections using Fiji (https://imagej.net/software/fiji/). In Figures 2E and 4D, only cells that imported Whi5 30 minutes after the start of imaging were included in the analysis. Maximum projections are shown throughout, except in Figure 4A-B, where sum projections are shown.

Quantification of GFP or mCherry fusion protein abundance was determined in background-subtracted 2D sum projections of whole-cell Z-stacks, with the nuclear area defined by Nup49-mCherry or by Nup60-mCherry. For Sac3 and Mtr2 quantifications (Figure 4A) G1 mother and daughter cells were quantified in G1, defined as the first 30-45 min after completion of anaphase and the absence of bud. For the *GAL1pr:sfGFP* reporter, the GFP mean fluorescence was determined (Figure 5). For the mother/daughter measurements (Figure 5C and 5E), mother/daughter pairs with individual nuclei in the moment of galactose shift were tracked and their fluorescence was measured at the indicated times. For the quantification of *GAL1pr:sfGFP* activation over time (Figure 5B and 5D), a custom Fiji macros segmented the nuclear signal (Nup60-mCherry) and, after manual correction of ROI, total fluorescence and mean fluorescence of the mCherry and GFP channels was automatically determined for all the individual cells.

Conventional epi-fluorescence microscopy was carried out with a Leica DM4000B wide-field microscope equipped with HCX PL APO 100X/1.40 OIL PH3 CS objective, image acquisition was performed with Hamamatsu ORCA-Flash4.0 LT digital CMOS camera with the help of Leica Application Suite X (LAS X) software. For DNA staining, cells were fixed for 5 min by addition of 70% ethanol and resuspended in 1 μg/ml DAPI (4′,6-diamidino-2-phenylindole). Budding index was scored manually using the transmitted light images and the cell-counter plug-in of Fiji.

### Western blotting

Approximately 10 ml of exponential growing cells (OD_600_=0.3-0.6) were collected, resuspended in 200 μl of 0.1 M NaOH and incubated for 5 min at room temperature. Cells were collected by centrifugation, resuspended in 50 μl of Laemmli buffer, and incubated for 5 min at 95 °C. Extracts were clarified by centrifugation and equivalent amounts of protein were resolved in an SDS-PAGE electrophoresis and transferred onto a nitrocellulose membrane. Membranes were blocked with milk powder 5% in TBS Tween 0.01% or FBS 10% in TBS Tween 0.1% (anti-AcLys) and incubated overnight with primary antibodies. Primary antibodies were anti-HA peroxidase 3F10 (Roche Diagnostics, Cat. No: 12013819001) diluted 1:5000, anti-GFP (Roche Diagnostics, Cat. No: 11814460001) diluted 1:5000, anti-GAPDH (Thermo Fisher Scientific Cat. No: MA5-15738) diluted 1:2000, anti-G-6-PDH diluted 1:20000 (Sigma, Cat. No: A9521) and anti AcLys diluted 1:1000 (Cell Signalling, Cat. No: 9681). Blots were developed with anti-mouse IgG and anti-rabbit IgG Horseradish Peroxidase conjugate (Thermo Fisher Scientific, Cat. No: 170-6516 or 31460 respectively) diluted 1:20000 using the Supersignal West Femto Maximum Sensitivity Substrate (Thermo Scientific). Bands were quantified with ImageQuant^TM^ LAS 4000 mini biomolecular imager (GE Healthcare). Uncropped western blots are shown in **Supplementary Figure S12.**

### Protein Immunoprecipitation assays

Approximately 100 ml of exponential growing cells (OD_600_=0.5-0.8) expressing Nup60-GFP were collected and resuspended in 300 μl of Lysis Buffer (50 mM Tris-HCl pH 8, 250 mM NaCl, 5 mM EDTA, 1% Triton X-100, 1 mM PMSF, 2.5mM Benzamidine, 0.1% (w/v) sodium deoxycholate) with Complete Mini protease inhibitor (Roche Diagnostics) and Histone deacetylase inhibitors (iHDAC) 5 μM Trichostatin A (Sigma Aldrich), 10 mM Nicotinamide (Sigma Aldrich) and 10 mM Sodium Butyrate (Sigma Aldrich). Cells were broken with vigorous shaking in the presence of glass beads, the cellular debris was removed and the supernatant was clarified by centrifugation at 13400×g for 5 min. 50 μl Dynabeads Protein G magnetic beads (Invitrogen) were incubated with anti-GFP antibody (Roche Diagnostics) for 20 min at room temperature and after washing with phosphate-buffered saline (PBS) containing Tween 0.02%, incubated with the cell extract for 2h at 4°C. After washing the beads 4 times with PBS Tween 0.02%, the immunoprecipitated Nup60 was eluted by boiling the beads for 5 min with 100 μl of Laemmli buffer 2X and analyzed by SDS-PAGE electrophoresis followed by western blot analysis.

### PolyA Fluorescence In Situ Hybridization (FISH)

Cells were grown in SC until exponential phase at 25 °C and the culture was divided in two to incubate half of the culture at 25 °C or 37 °C or in SC or SC-gal during the indicated times. Approximately 10 ml of exponential cultures (OD_600_ = 0.5-1) were fixed with 4% (v/v) formaldehyde (Sigma-Aldrich) during 15 min at room temperature or 37 °C. Cells were collected and resuspended in 0.1 M potassium phosphate (KPi) pH = 6.4/4% formaldehyde and fixated for 1h in agitation at room temperature. The fixation agent was removed by washing two times with 0.1 M KPi pH 6.4. Cells were washed one time with ice-cold washing buffer (0.1 M KPi (pH = 6.4)/1.2 M sorbitol), resuspended in 200 μl of ice-cold washing buffer and subsequently cell wall was digested by incubating 100 μl of cells with 0.4 mg/ml of Zymolyase 100T (SEIKAGAKU CORPORATION) at 30 °C for 5-15 min. Partially spheroplasted cells were recovered by centrifugation (3000 g for 5 min), washed one time with washing buffer and resuspended in 30 μl 0.1 M KPi (pH = 6.4)/1.2 M sorbitol. Samples were then applied on Teflon slides with wells previously coated with poly-L-lysine. Non-adhering cells were removed by aspiration, cells were rehydrated with 2X SSC (0.15M NaCl and 0.015M sodium citrate) and incubated with prehybridization buffer (formamide 50%, dextran sulfate 10%, 125μg/ml of Escherichia coli tRNA, 4X SSC, 1X Denhardt’s solution and 500 μg/ml herring sperm DNA) for 1h at 37 °C in in a humid chamber. The hybridization was incubated overnight at 37 °C in a humid chamber with 20 μl of prehybridization buffer supplemented with 1 μM of Cy3-end-labeled oligo(dT), 1 mM DTT and RNasin (Promega) 4U/ml. After hybridization, slides were washed with 2X SSC and 1X SSC at room temperature for 5 min and subsequently cells were incubated for 1 min with 2 μl of DAPI 2.5 mg/ml. Cells were washed twice with 1X SSC, washed with 0.5X SSC, air dried and mounted with 50% glycerol. Detection of Cy3-oligo(dT) and DAPI was performed using a Leica DM4000B fluorescence microscope.

### Flow cytometry

DNA content analysis was performed according to the protocol from (Rosebrock, 2017). Briefly, cells were fixed in 2 volumes of 100% ethanol (overnight -20℃), rehydrated with PBS and digested with proteinase K (20 μg/ml, Thermo Scientific) and RNAse A (10 μg/ml) for 2h at 50℃ in the presence of SYTOX Green (0.5 μM, Invitrogen). Flow cytometry was performed on a LSRII (BD) instrument and the data was analysed with FlowJo software. Gating strategies are shown in **Supplementary Figure S13.**

### Gene expression analysis (RT-qPCR)

Samples for gene expression analysis containing approximately 4x10^8^ cells were frozen in liquid nitrogen and stored at -80℃. RiboPure-Yeast (Invitrogen) kit was used to isolate total RNA according to the manufacturer’s protocol. Briefly, cells were broken in a Disruptor Genie (Scientific Industries) with Zirconia Beads in Lysis Buffer with one volume of Phenol:Chloroform. The lysate was centrifuged at 16,100 xg for 10 min, and the aqueous phase was separated and mixed with corresponding amounts of 100% ethanol and Binding Buffer. The resulting solution was drawn through the column filter which was further washed with Wash Solutions. RNA was eluted from the filter with 2 x 50 µl of Elution Buffer and 1 x 50 µl with DEPC-treated Molecular Biology Grade water (Merck). RNA samples were quantified with NanoDrop 2000 Spectrophotometer (Thermo Scientific). 2.5 μg of RNA were incubated with ezDNase and cDNA was obtained with SuperScript IV VILO Master Mix (both Invitrogen) following the manufacturer instructions. cDNA was analyzed in triplicate by quantitative RT-PCR with LightCycler 480 SYBR Green I Master mix (Roche) using the Roche LightCycler 480 II instrument. mRNA levels of genes of interest were quantified relative to *ACT1* mRNA by the ΔCt method.

### Statistical methods and reproducibility

Graphs and statistical analysis (Two-tailed unpaired t-test and Log-rank Mantel-Cox test) were performed with GraphPad Prism software and R. For all the strains used in this work, at least 2 independent clones have been tested with similar results. All the data has been obtained from at least 2 independent biological replicates.

## References

Aksenova, V., Smith, A., Lee, H., Bhat, P., Esnault, C., Chen, S., et al. (2020). Nucleoporin TPR is an integral component of the TREX-2 mRNA export pathway. Nat. Commun. 11, 4577.

Allard, S., Utley, R. T., Savard, J., Clarke, A., Grant, P., Brandl, C. J., et al. (1999). NuA4, an essential transcription adaptor/histone H4 acetyltransferase complex containing Esa1p and the ATM-related cofactor Tra1p. EMBO J. 18, 5108–5119.

Baptista, T., Grünberg, S., Minoungou, N., Koster, M. J. E., Timmers, H. T. M., Hahn, S., et al. (2017). SAGA Is a General Cofactor for RNA Polymerase II Transcription. Mol. Cell 68, 130–143.e5.

Bertoli, C., Skotheim, J. M., and de Bruin, R. A. M. (2013). Control of cell cycle transcription during G1 and S phases. Nat. Rev. Mol. Cell Biol. 14, 518–528.

Brickner, D. G., Randise-Hinchliff, C., Lebrun Corbin, M., Liang, J. M., Kim, S., Sump, B., et al. (2019). The Role of Transcription Factors and Nuclear Pore Proteins in Controlling the Spatial Organization of the Yeast Genome. Dev. Cell 49, 936–947.e4.

Bruzzone, M. J., Grünberg, S., Kubik, S., Zentner, G. E., and Shore, D. (2018). Distinct patterns of histone acetyltransferase and Mediator deployment at yeast protein-coding genes. Genes Dev. 32, 1252–1265.

Cabal, G. G., Genovesio, A., Rodriguez-Navarro, S., Zimmer, C., Gadal, O., Lesne, A., et al. (2006). SAGA interacting factors confine sub-diffusion of transcribed genes to the nuclear envelope. Nature 441, 770–773.

Casolari, J. M., Brown, C. R., Komili, S., West, J., Hieronymus, H., and Silver, P. A. (2004). Genome-wide localization of the nuclear transport machinery couples transcriptional status and nuclear organization. Cell 117, 427–439.

Charvin, G., Oikonomou, C., Siggia, E. D., and Cross, F. R. (2010). Origin of irreversibility of cell cycle start in budding yeast. PLoS Biol. 8, e1000284.

Chen, P. B., Hung, J.-H., Hickman, T. L., Coles, A. H., Carey, J. F., Weng, Z., et al. (2013). Hdac6 regulates Tip60-p400 function in stem cells. Elife 2, e01557.

Chen, Y., Zhao, G., Zahumensky, J., Honey, S., and Futcher, B. (2020). Differential Scaling of Gene Expression with Cell Size May Explain Size Control in Budding Yeast. Mol. Cell 78, 359–370.e6.

Choudhary, C., Kumar, C., Gnad, F., Nielsen, M. L., Rehman, M., Walther, T. C., et al. (2009). Lysine acetylation targets protein complexes and co-regulates major cellular functions. Science 325, 834–840.

Clarke, A. S., Lowell, J. E., Jacobson, S. J., and Pillus, L. (1999). Esa1p is an essential histone acetyltransferase required for cell cycle progression. Mol. Cell. Biol. 19, 2515–2526.

Costanzo, M., Nishikawa, J. L., Tang, X., Millman, J. S., Schub, O., Breitkreuz, K., et al. (2004). CDK activity antagonizes Whi5, an inhibitor of G1/S transcription in yeast. Cell 117, 899–913.

de Bruin, R. A. M., Mcdonald, W. H., Kalashnikova, T. I., Yates, J., and Wittenberg, C. (2004). Cln3 activates G1-specific transcription via phosphorylation of the SBF bound repressor Whi5. Cell 117, 887–898.

Dieppois, G., Iglesias, N., and Stutz, F. (2006). Cotranscriptional recruitment to the mRNA export receptor Mex67p contributes to nuclear pore anchoring of activated genes. Mol. Cell. Biol. 26, 7858–7870.

Di Talia, S., Skotheim, J. M., Bean, J. M., Siggia, E. D., and Cross, F. R. (2007). The effects of molecular noise and size control on variability in the budding yeast cell cycle. Nature 448, 947–951.

Di Talia, S., Wang, H., Skotheim, J. M., Rosebrock, A. P., Futcher, B., and Cross, F. R. (2009). Daughter-specific transcription factors regulate cell size control in budding yeast. PLoS Biol. 7, e1000221.

Downey, M., Johnson, J. R., Davey, N. E., Newton, B. W., Johnson, T. L., Galaang, S., et al. (2015). Acetylome profiling reveals overlap in the regulation of diverse processes by sirtuins, gcn5, and esa1. Mol. Cell. Proteomics 14, 162–176.

Doyon, Y., and Côté, J. (2004). The highly conserved and multifunctional NuA4 HAT complex. Curr. Opin. Genet. Dev. 14, 147–154.

Estruch, F., Peiró-Chova, L., Gómez-Navarro, N., Durbán, J., Hodge, C., Del Olmo, M., et al. (2009). A genetic screen in Saccharomyces cerevisiae identifies new genes that interact with mex67-5, a temperature-sensitive allele of the gene encoding the mRNA export receptor. Mol. Genet. Genomics 281, 125–134.

Fischer, T., Strässer, K., Rácz, A., Rodriguez-Navarro, S., Oppizzi, M., Ihrig, P., et al. (2002). The mRNA export machinery requires the novel Sac3p-Thp1p complex to dock at the nucleoplasmic entrance of the nuclear pores. EMBO J. 21, 5843–5852.

Frolov, M. V., and Dyson, N. J. (2004). Molecular mechanisms of E2F-dependent activation and pRB-mediated repression. J. Cell Sci. 117, 2173–2181.

Gallego, O., Specht, T., Brach, T., Kumar, A., Gavin, A.-C., and Kaksonen, M. (2013). Detection and characterization of protein interactions in vivo by a simple live-cell imaging method. PLoS One 8, e62195.

Gelperin, D. M., White, M. A., Wilkinson, M. L., Kon, Y., Kung, L. A., Wise, K. J., et al. (2005). Biochemical and genetic analysis of the yeast proteome with a movable ORF collection. Genes Dev. 19, 2816–2826.

Gomar-Alba, M., and Mendoza, M. (2019). Modulation of Cell Identity by Modification of Nuclear Pore Complexes. Front. Genet. 10, 1301.

Goulev, Y., Matifas, A., Heyer, V., Reina-San-Martin, B., and Charvin, G. (2019). COSPLAY: An expandable toolbox for combinatorial and swift generation of expression plasmids in yeast. PLoS One 14, e0220694.

Hampoelz, B., Andres-Pons, A., Kastritis, P., and Beck, M. (2019). Structure and Assembly of the Nuclear Pore Complex. Annu. Rev. Biophys. 48, 515–536.

Helmlinger, D., and Tora, L. (2017). Sharing the SAGA. Trends Biochem. Sci. 42, 850–861.

Henriksen, P., Wagner, S. A., Weinert, B. T., Sharma, S., Bacinskaja, G., Rehman, M., et al. (2012). Proteome-wide analysis of lysine acetylation suggests its broad regulatory scope in Saccharomyces cerevisiae. Mol. Cell. Proteomics 11, 1510–1522.

Huang, D., Kaluarachchi, S., van Dyk, D., Friesen, H., Sopko, R., Ye, W., et al. (2009). Dual regulation by pairs of cyclin-dependent protein kinases and histone deacetylases controls G1 transcription in budding yeast. PLoS Biol. 7, e1000188.

Ibarra, A., and Hetzer, M. W. (2015). Nuclear pore proteins and the control of genome functions. Genes Dev. 29, 337–349.

Jani, D., Valkov, E., and Stewart, M. (2014). Structural basis for binding the TREX2 complex to nuclear pores, GAL1 localisation and mRNA export. Nucleic Acids Res. 42, 6686–6697.

Janke, C., Magiera, M. M., Rathfelder, N., Taxis, C., Reber, S., Maekawa, H., et al. (2004). A versatile toolbox for PCR-based tagging of yeast genes: new fluorescent proteins, more markers and promoter substitution cassettes. Yeast 21, 947–962.

Jones, G. M., Stalker, J., Humphray, S., West, A., Cox, T., Rogers, J., et al. (2008). A systematic library for comprehensive overexpression screens in Saccharomyces cerevisiae. Nat. Methods 5, 239–241.

Kadowaki, T., Hitomi, M., Chen, S., and Tartakoff, A. M. (1994). Nuclear mRNA accumulation causes nucleolar fragmentation in yeast mtr2 mutant. Mol. Biol. Cell 5, 1253–1263.

Kaluarachchi Duffy, S., Friesen, H., Baryshnikova, A., Lambert, J.-P., Chong, Y. T., Figeys, D., et al. (2012). Exploring the yeast acetylome using functional genomics. Cell 149, 936–948.

Kehat, I., Accornero, F., Aronow, B. J., and Molkentin, J. D. (2011). Modulation of chromatin position and gene expression by HDAC4 interaction with nucleoporins. J. Cell Biol. 193, 21–29.

Kishkevich, A., Cooke, S. L., Harris, M. R. A., and de Bruin, R. A. M. (2019). Gcn5 and Rpd3 have a limited role in the regulation of cell cycle transcripts during the G1 and S phases in Saccharomyces cerevisiae. Sci. Rep. 9, 1–9.

Knockenhauer, K. E., and Schwartz, T. U. (2016). The Nuclear Pore Complex as a Flexible and Dynamic Gate. Cell 164, 1162–1171.

Kumar, A., Sharma, P., Gomar-Alba, M., Shcheprova, Z., Daulny, A., Sanmartín, T., et al. (2018). Daughter-cell-specific modulation of nuclear pore complexes controls cell cycle entry during asymmetric division. Nat. Cell Biol. 20, 432–442.

Kurshakova, M. M., Krasnov, A. N., Kopytova, D. V., Shidlovskii, Y. V., Nikolenko, J. V., Nabirochkina, E. N., et al. (2007). SAGA and a novel Drosophila export complex anchor efficient transcription and mRNA export to NPC. EMBO J. 26, 4956–4965.

Lee, K. K., and Workman, J. L. (2007). Histone acetyltransferase complexes: one size doesn’t fit all. Nat. Rev. Mol. Cell Biol. 8, 284–295.

Light, W. H., Brickner, D. G., Brand, V. R., and Brickner, J. H. (2010). Interaction of a DNA zip code with the nuclear pore complex promotes H2A.Z incorporation and INO1 transcriptional memory. Mol. Cell 40, 112–125.

Li, Z., Vizeacoumar, F. J., Bahr, S., Li, J., Warringer, J., Vizeacoumar, F. S., et al. (2011). Systematic exploration of essential yeast gene function with temperature-sensitive mutants. Nat. Biotechnol. 29, 361–367.

Longtine, M. S., McKenzie, A., Demarini, D. J., Shah, N. G., Wach, A., Brachat, A., et al. (1998). Additional modules for versatile and economical PCR-based gene deletion and modification in Saccharomyces cerevisiae. Yeast 14, 953–961.

Narita, T., Weinert, B. T., and Choudhary, C. (2019). Functions and mechanisms of non-histone protein acetylation. Nat. Rev. Mol. Cell Biol. 20, 156–174.

Raices, M., and D’Angelo, M. A. (2021). Structure, Maintenance, and Regulation of Nuclear Pore Complexes: The Gatekeepers of the Eukaryotic Genome. Cold Spring Harb. Perspect. Biol. doi:10.1101/cshperspect.a040691.

Rosebrock, A. P. (2017). Analysis of the Budding Yeast Cell Cycle by Flow Cytometry. Cold Spring Harb. Protoc. 2017. doi:10.1101/pdb.prot088740.

Schmoller, K. M., Turner, J. J., Kõivomägi, M., and Skotheim, J. M. (2015). Dilution of the cell cycle inhibitor Whi5 controls budding-yeast cell size. Nature 526, 268–272.

Schneider, M., Hellerschmied, D., Schubert, T., Amlacher, S., Vinayachandran, V., Reja, R., et al. (2015). The Nuclear Pore-Associated TREX-2 Complex Employs Mediator to Regulate Gene Expression. Cell 162, 1016–1028.

Schneiter, R., Kadowaki, T., and Tartakoff, A. M. (1995). mRNA transport in yeast: time to reinvestigate the functions of the nucleolus. Mol. Biol. Cell 6, 357–370.

Skotheim, J. M., Di Talia, S., Siggia, E. D., and Cross, F. R. (2008). Positive feedback of G1 cyclins ensures coherent cell cycle entry. Nature 454, 291–296.

Sood, V., and Brickner, J. H. (2014). Nuclear pore interactions with the genome. Curr. Opin. Genet. Dev. 25, 43–49.

Sterner, D. E., and Berger, S. L. (2000). Acetylation of histones and transcription-related factors. Microbiol. Mol. Biol. Rev. 64, 435–459.

Strässer, K., Bassler, J., and Hurt, E. (2000). Binding of the Mex67p/Mtr2p heterodimer to FXFG, GLFG, and FG repeat nucleoporins is essential for nuclear mRNA export. J. Cell Biol. 150, 695–706.

Strawn, L. A., Shen, T., and Wente, S. R. (2001). The GLFG regions of Nup116p and Nup100p serve as binding sites for both Kap95p and Mex67p at the nuclear pore complex. J. Biol. Chem. 276, 6445–6452.

Takahata, S., Yu, Y., and Stillman, D. J. (2009). The E2F functional analogue SBF recruits the Rpd3(L) HDAC, via Whi5 and Stb1, and the FACT chromatin reorganizer, to yeast G1 cyclin promoters. EMBO J. 28, 3378–3389.

Texari, L., Dieppois, G., Vinciguerra, P., Contreras, M. P., Groner, A., Letourneau, A., et al. (2013). The nuclear pore regulates GAL1 gene transcription by controlling the localization of the SUMO protease Ulp1. Mol. Cell 51, 807–818.

Turner, J. J., Ewald, J. C., and Skotheim, J. M. (2012). Cell Size Control in Yeast. Curr. Biol. 22, R350–R359.

Umlauf, D., Bonnet, J., Waharte, F., Fournier, M., Stierle, M., Fischer, B., et al. (2013). The human TREX-2 complex is stably associated with the nuclear pore basket. J. Cell Sci. 126, 2656–2667.

Veening, J.-W., Smits, W. K., and Kuipers, O. P. (2008). Bistability, epigenetics, and bet-hedging in bacteria. Annu. Rev. Microbiol. 62, 193–210.

Wang, H., Carey, L. B., Cai, Y., Wijnen, H., and Futcher, B. (2009). Recruitment of Cln3 cyclin to promoters controls cell cycle entry via histone deacetylase and other targets. PLoS Biol. 7, e1000189.

Wickramasinghe, V. O., Andrews, R., Ellis, P., Langford, C., Gurdon, J. B., Stewart, M., et al. (2014). Selective nuclear export of specific classes of mRNA from mammalian nuclei is promoted by GANP. Nucleic Acids Research 42, 5059–5071. doi:10.1093/nar/gku095.

Wickramasinghe, V. O., McMurtrie, P. I. A., Mills, A. D., Takei, Y., Penrhyn-Lowe, S., Amagase, Y., et al. (2010). mRNA export from mammalian cell nuclei is dependent on GANP. Curr. Biol. 20, 25–31.

