## Supplementary Figures for "Nuclear Pore Complex Acetylation Regulates mRNA Export and Cell Cycle Commitment in Budding Yeast"

Mercè Gomar-Alba *et al*.

**Supplementary Figures S1-S13 and Table 1**

**A**

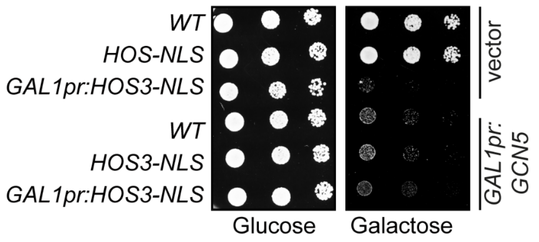


**B**

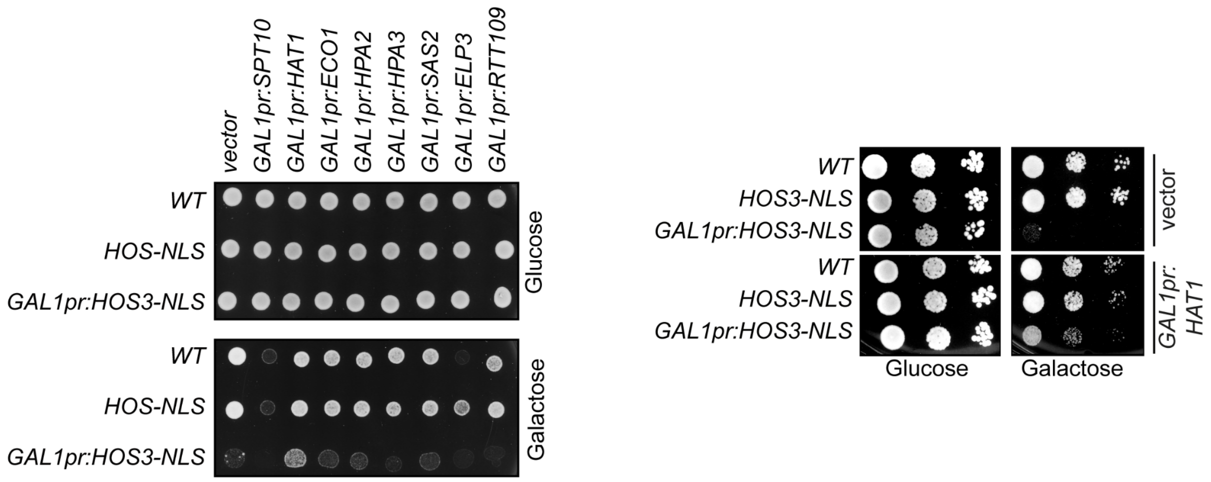


**Supplementary Figure S1**. **Effect of KAT overexpression in Hos3-NLS dependent growth inhibition. (A)** Overexpression of the KAT Gcn5 is toxic. 10-fold serial dilutions of wild-type (*WT*), *HOS3-NLS-GFP* and *GAL1pr:HOS3-NLS-GFP* (single copy at the endogenous locus) transformed with an empty vector or the *GAL1pr:GCN5-HA* plasmid, were spotted onto SC-Glu and SC-Gal medium and incubated at 25 °C for 3 days. **(B)** Role of KAT overexpression in cell viability and ability to rescue growth in the presence of overexpressed *HOS3-NLS*. (*Left*) Exponential cultures of the indicated strains transformed with an empty vector or plasmids overexpressing the KATs Spt10, Eco1, Hpa2, Hpa3, Sas2, Elp3 or Rtt109, were spotted onto SC-Glu and SC-Gal medium and incubated at 25 °C for 3 days. (*Right*) The effect of *HAT1* overexpression was also assessed in 10-fold serial dilutions as in (A).

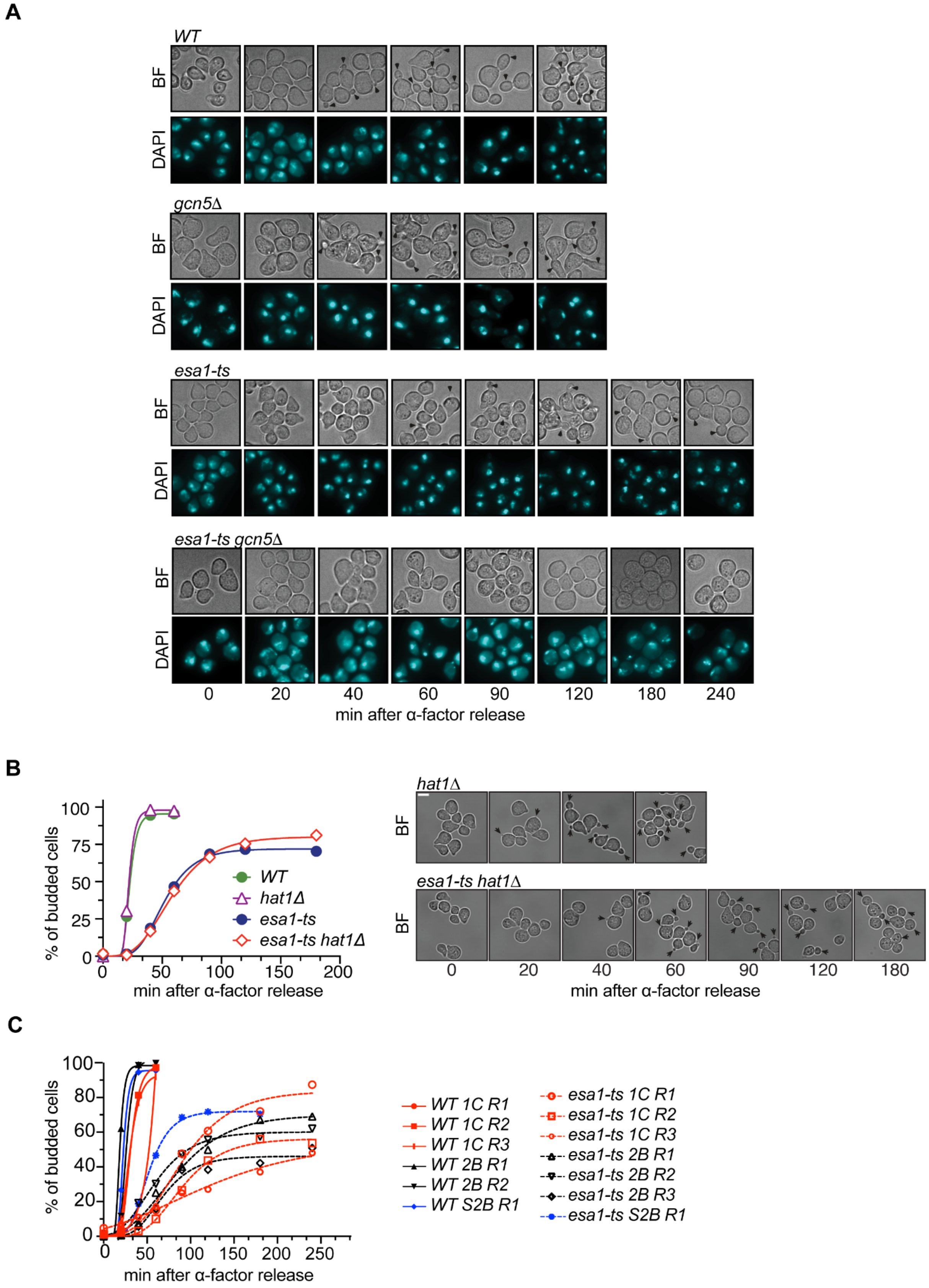


**Supplementary Figure S2**. **Role of *ESA1*, *GCN5* and *HAT1* in Start.** **(A)** *esa1-ts* and *gcn5Δ esa1-ts* mutants have bud emergence defects. Cells of the indicated strains were arrested in G1 by treatment with α-factor for 2.5 h at 25 ºC, shifted to 37 ºC for 1 h and released from the G1 arrest at 37 ºC. Cells were fixed at the indicated times and the presence of buds (arrowheads) was assessed by microscopy. **(B)** *HAT1* is not involved in Start. Bright field images of cells of the indicated strains, treated as in (A) and scored at the indicated times after α-factor washout. At least 200 cells were scored for each strain and time point. Arrowheads point to cell buds. Scale bar, 5 μm. **(C)** Variability of *esa1-ts* budding defects. Quantification of the budding index is shown for all replicates (R) shown in figures 1C, 2C and S2B. BF, brightfield; DAPI, nuclear stain.

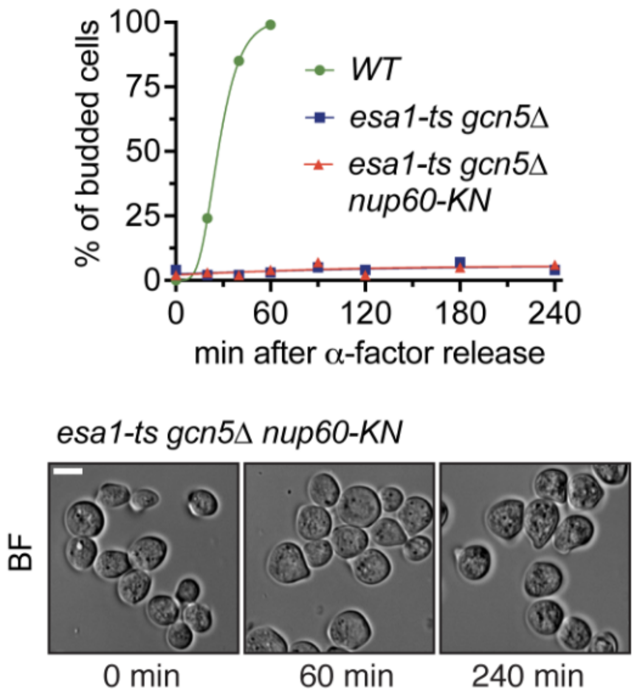


**Supplementary Figure S3**. **Acetyl-mimic *nup60-KN* mutation does not rescue budding in *esa1-ts gcn5Δ* double mutant cells.** Cells of the indicated strains were arrested in G1 by treatment with α-factor for 2.5 h at 25 ºC, shifted to 37 ºC for 1 h and released from the G1 arrest at 37 ºC. Cells were fixed at the indicated times and the presence of buds was assessed by microscopy. At least 200 cells were scored for each strain and time point. Bright field images of the indicated strains at the indicated times after the α-factor washout. Arrowheads point to cell buds. Scale bar, 5 μm.

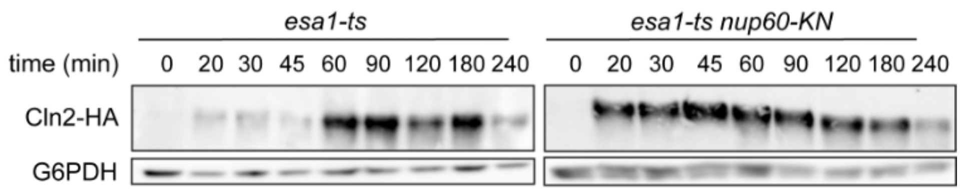


**Supplementary Figure S4.** ***nup60-KN*** **mutation partially rescues the delay in synthesis of the G1/S cyclin Cln2 in** ***esa1-ts*** **cells.** Independent biological replicate of the experiment shown in Figure 2D. Cells of the indicated strains were arrested in G1 by treatment with α-factor for 2.5 h at 25 ºC, shifted to 37 ºC for 1 h and released from the G1 arrest at 37 ºC. Samples for total protein extracts were collected at the indicated times after α-factor washout and the amount of Cln2-HA protein was assessed by western blot. G6PDH was used as loading control.

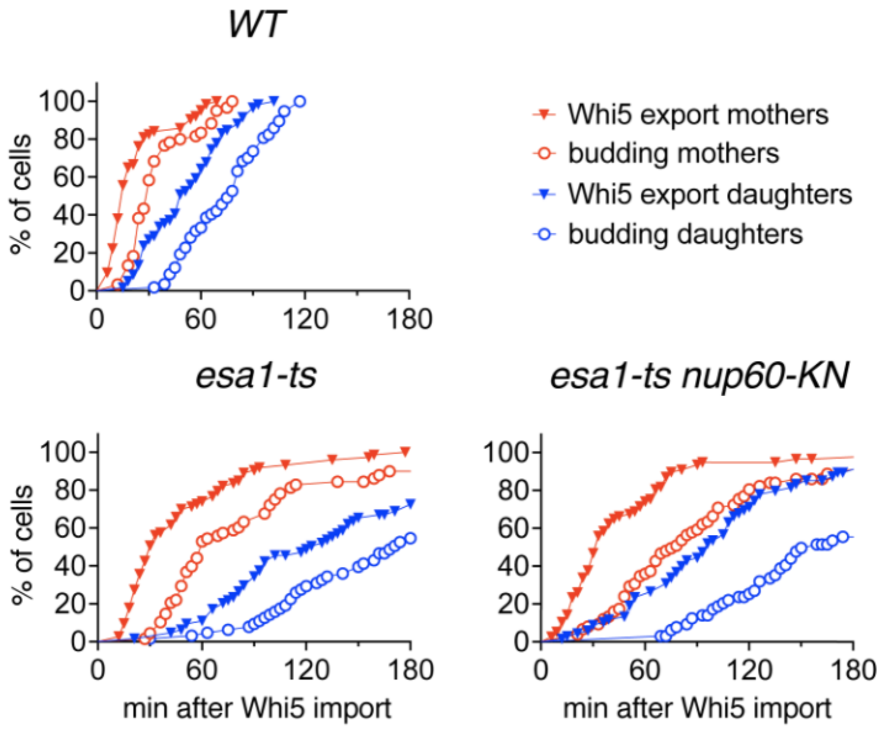


**Supplementary Figure S5. Whi5-mCherry nuclear export and budding for cells in Figure 2E**. Whi5-mCherry nuclear export was scored in the fluorescence channel, and budding was scored in bright-field images (maximum projections of 3 z-confocal slices spaced 0.5 µm).

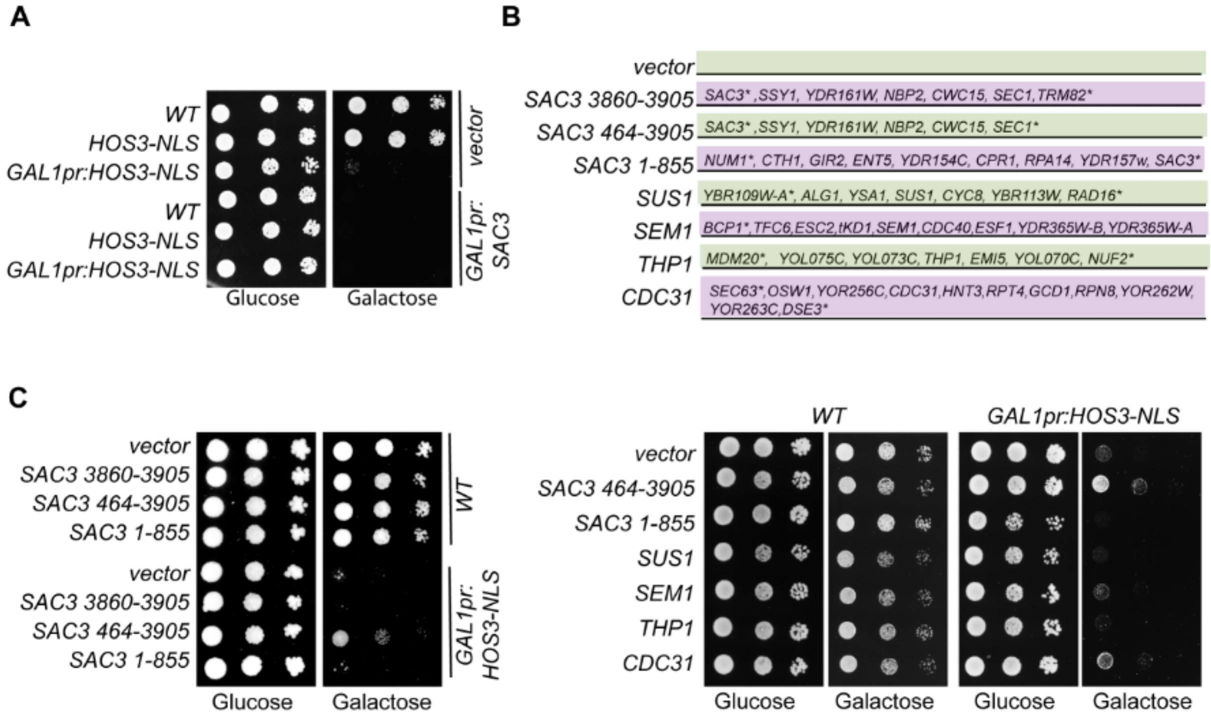


**Supplementary Figure S6. Overexpression of the TREX-2 complex component Sac3 rescues the toxicity of *HOS3-NLS* overexpression. (A)** Toxicity of full-length *SAC3* overexpression. 10-fold serial dilutions of wild-type (*WT*), *HOS3-NLS-GFP* and *GAL1pr:HOS3-NLS-GFP* cultures transformed with an empty vector or the *GAL1pr:SAC3-HA* plasmid were spotted onto SC-Glu and SC-Gal medium and incubated at 25 °C for 3 days. **(B)** List of high-copy (2µ) plasmids from a tiling genome library (Jones et al., 2008) containing TREX-2 complex genes (*SAC3*, *SUS1*, *CDC31*, *SEM1* and *THP1*) together with neighboring genes. Asterisks (*) indicate that the corresponding ORFs are incomplete. Nucleotides of *SAC3* (full length 3905 nt) included in each plasmid are indicated. **(C)** A high-copy plasmid containing *SAC3(464-3905)* is not toxic and relieves the toxicity of *HOS3-NLS* over-expression. 10-fold serial dilutions of wild-type (*WT*) and *GAL1pr:HOS3-NLS-GFP* cultures, transformed with an empty vector or the indicated multicopy plasmids, were spotted onto SC-Glu and SC-Gal medium and incubated at 25 °C for 3 days. Note that *SAC3(464-3905)* rescues growth of *GAL1pr:HOS3-NLS* but that the overlapping plasmid *SAC3(3860-3905)*, lacking all of *SAC3* ORF but 45 nucleotides at its 3’, does not. The “vector” and *SAC3(464-3905)* sections of the left image are also shown in Figure 3C for simplicity.

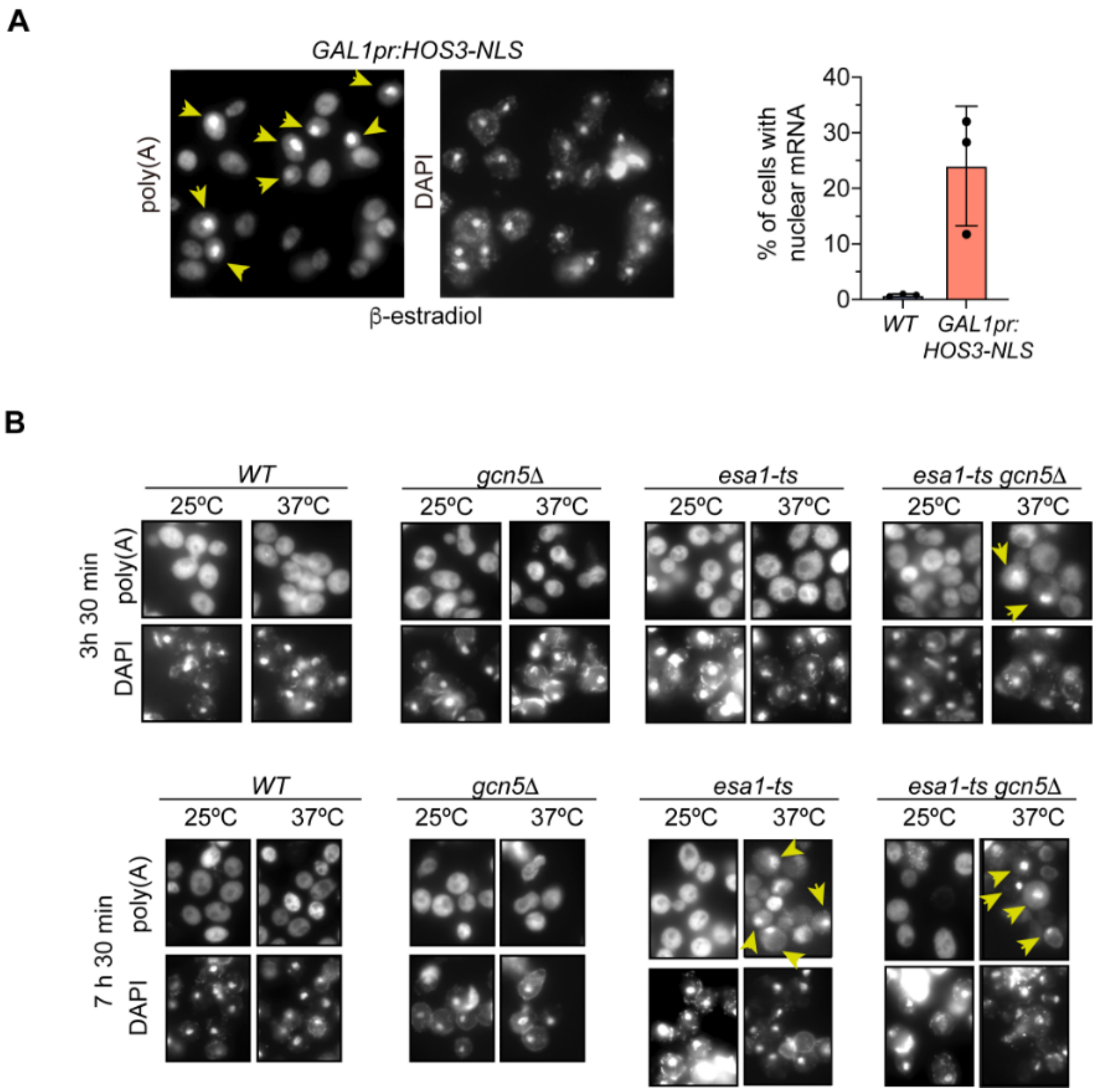


**Supplementary Figure S7. Hos3-NLS overexpression or depletion of Esa1 impairs export of poly(A) RNA. (A)** Overexpression of Hos3-NLS promotes nuclear accumulation of mRNA. Cultures of the indicated strains were treated with β-estradiol (90 nM) to induce Hos3-NLS. After induction overnight, cells were fixed and FISH was performed using a Cy3-Oligo(dT) probe. Arrows point to polyadenylated RNA in the nucleus, which was visualized by DAPI staining (*left*). The fraction of cells with nuclear mRNA accumulation was determined for the indicated strains and conditions (*right*). **(B)** Representative images of cells processed for poly(A) FISH as in (A) after incubation in the indicated conditions. Associated with Figure 3E.

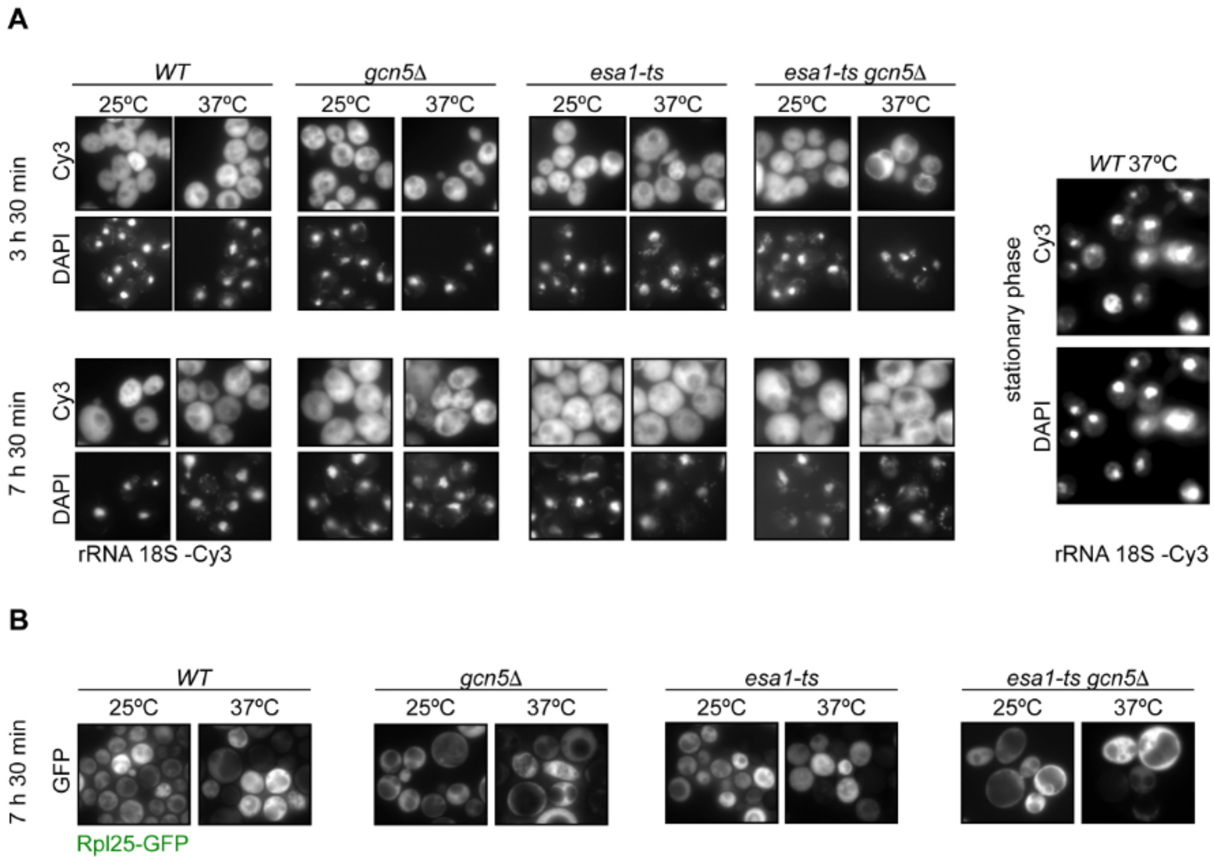


**Supplementary Figure S8. Depletion of Esa1 or Gcn5 does not affect export of rRNA. (A)** Wild-type, *gcn5*Δ, *esa1-ts* and *gcn5*Δ *esa1-ts* cultures were incubated at 25ºC or 37ºC at the indicated times. In all cases, cells were fixed and in situ hybridization was performed using Cy3-TXGTTCCTCGTTAAGGXATTTACATTGTACTXCC-Cy3 to target 18S rRNA and monitor ribosomal 40S subunit nucleo/cytoplasmic distribution. DNA was visualized by DAPI staining. Cells from stationary cultures exposed to heat shock during 4h were used as positive control for nuclear accumulation of 18S rRNA. **(B)** Wild-type, *gcn5*Δ, *esa1-ts* and *gcn5*Δ *esa1-ts* cultures transformed with an Rpl25-GFP plasmid as a reporter for ribosomal 60S subunit nucleo/cytoplasmic distribution were incubated at 25ºC or 37ºC for 7h and 30 min and imaged at the indicated conditions using fluorescence microscopy.

**A**

**B**

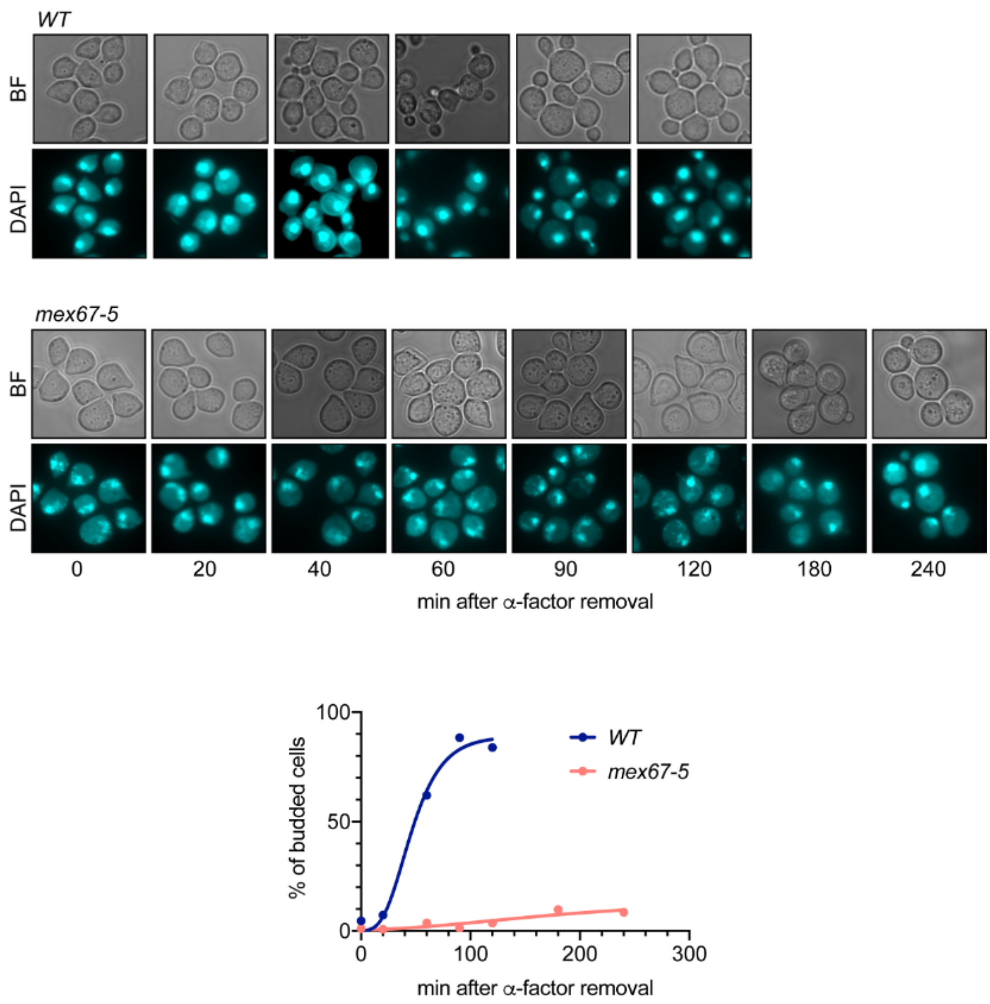


**Supplementary Figure S9. *mex67-5* thermosensitive mutant has bud emerge defects. (A)** Bright field (BF) images of wild type (*WT*) and *mex67-5* cells at the indicated times after the α-factor washout. Cells were arrested in G1 by treatment with α-factor for 2.5 h at 25 ºC, shifted to 37 ºC for 1 h and released from the G1 arrest at 37 ºC. The DNA was visualized by DAPI staining. (**B)** Cells were fixed at the indicated times and the presence of buds was assessed by microscopy. At least 200 cells were scored for each strain and time point.

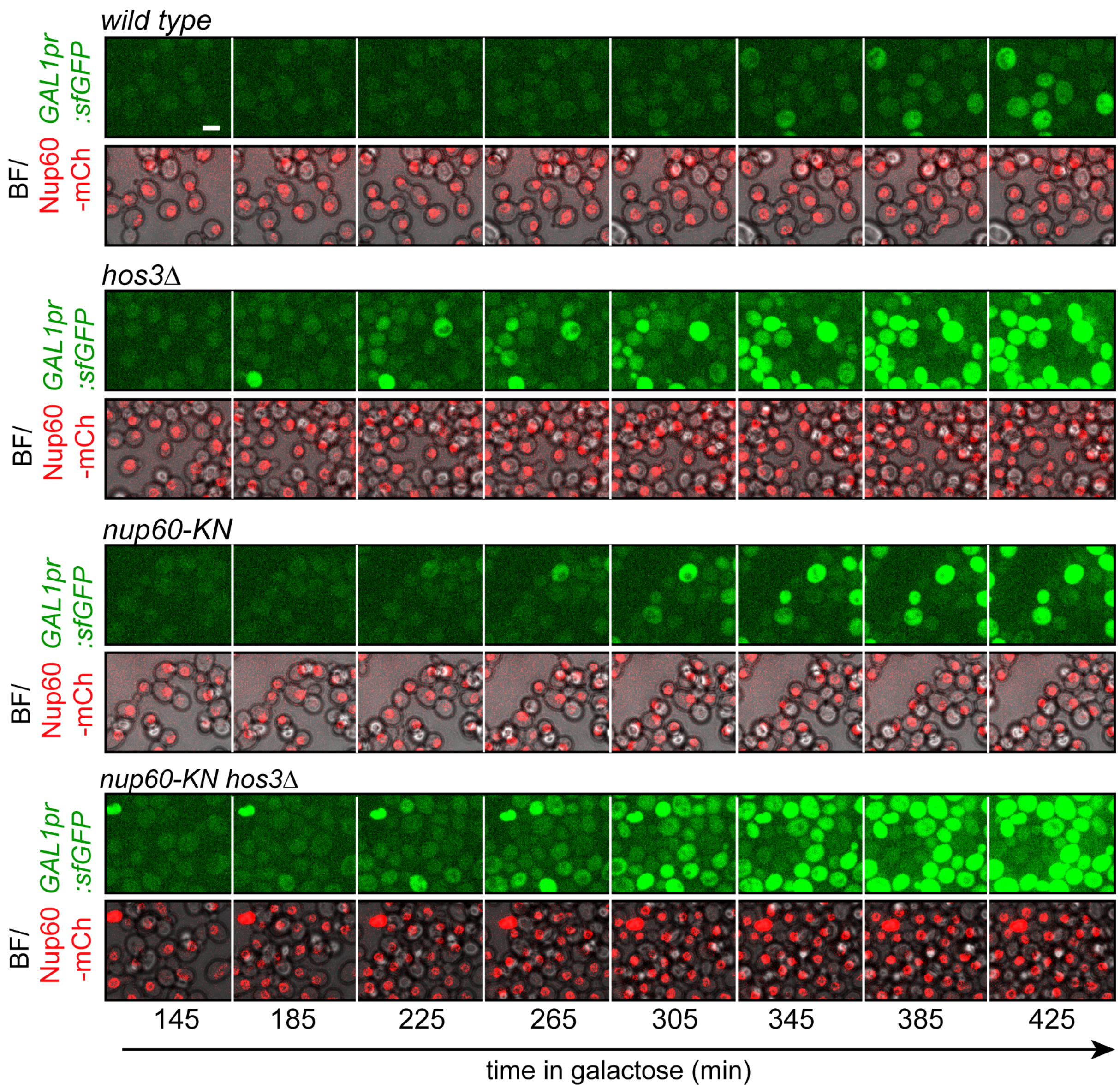


**Supplementary Figure S10.** Time Lapse microscopy of *WT, hos3*Δ*, nup60-KN hos3*Δ and *nup60-KN* cells expressing *GAL1pr:sfGFP* and Nup60-mCherry at the indicated times of galactose induction. Scale bar, 4 µm.

A

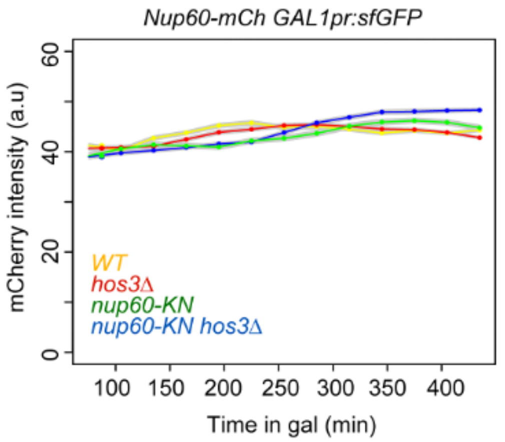


B

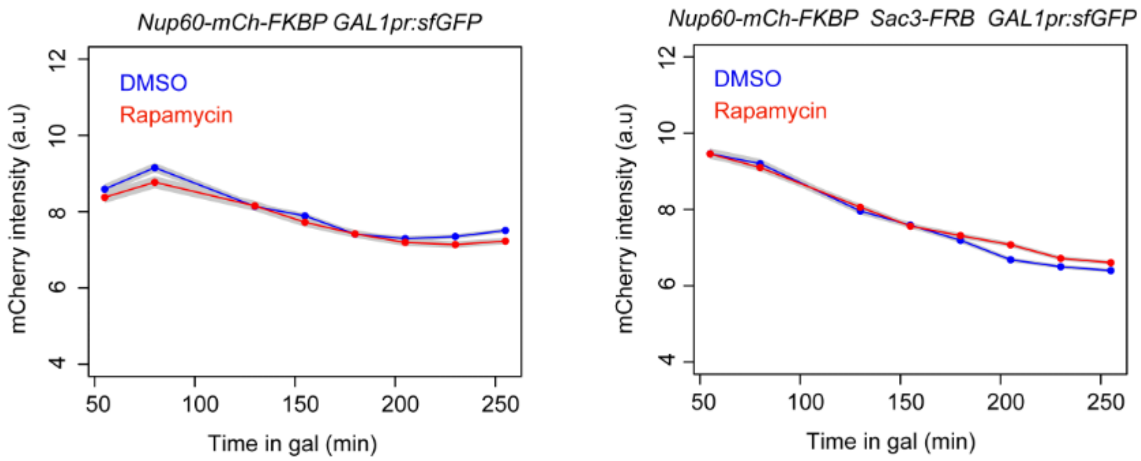


**Supplementary Figure S11. Nup60 protein levels upon galactose induction of *GAL1pr:sfGFP* are not changed by the acetyl-mimic allele of Nup60 (*nup60-KN*) or Hos3 and Sac3 anchoring to NPCs. (A)** Cultures of *WT*, *hos3Δ, nup60-KN* and *nup60-KN hos3Δ* were shifted to galactose and imaged by Time Lapse microscopy to monitor *Nup60-mCherry* fluorescence during 7 hours of galactose induction of *GAL1pr:sfGFP* expression. Nuclear fluorescence was scored by segmentation of the nuclear area in the mCherry channel and total fluorescence of Nup60-mCherry was quantified as in Figure 5B . At least 200 cells were scored for each strain and time point. Shaded areas indicate the SEM. **(B)** Cells expressing either *Nup60-mCherry-FKBP GAL1pr:sfGFP* or *Nup60-mCherry-FKBP Sac3-FRB GAL1pr:sfGFP* were incubated with rapamycin (RAPA) for FRB-FKBP heterodimerization or DMSO as control. Cells were imaged upon rapamycin and galactose addition and the *Nup60-mCherry* fluorescence over time was monitored as in A.

**Figure 2A**

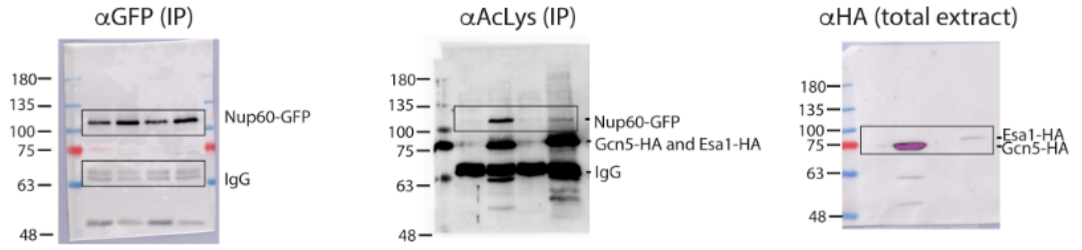


**Figure 2D**

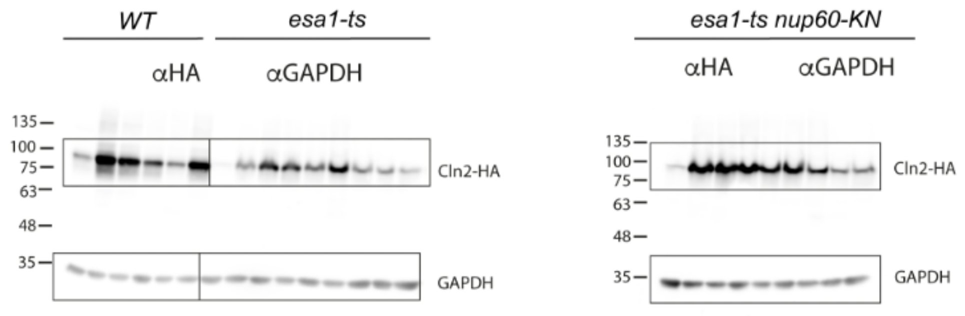


**Supplementary Figure S4**

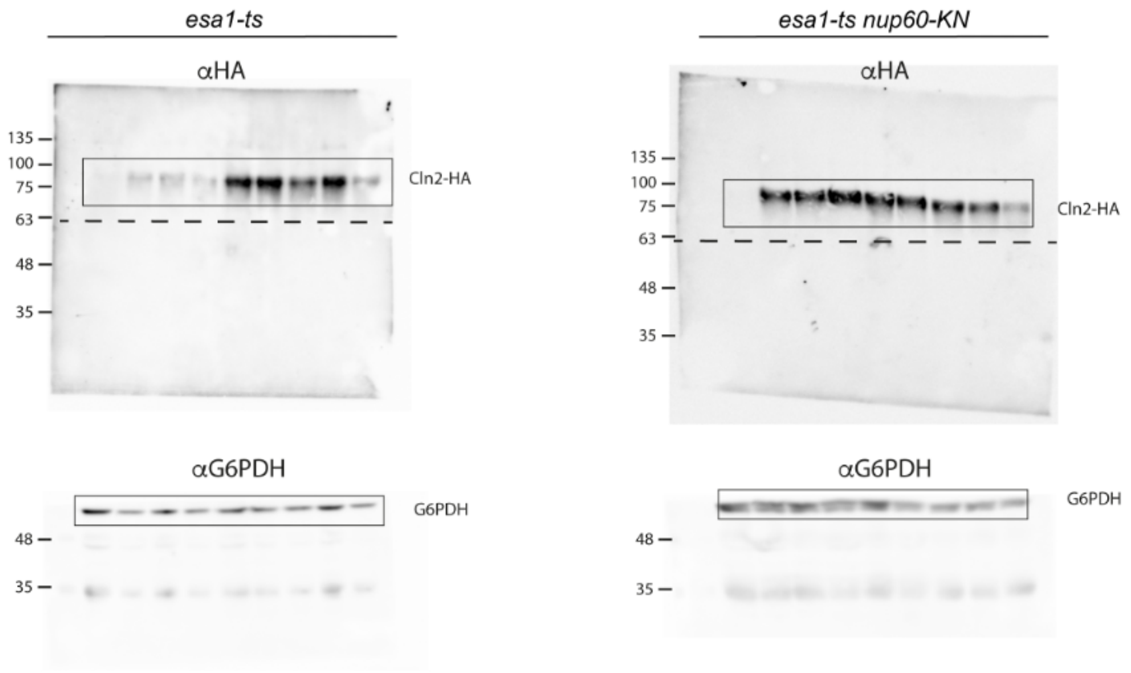


**Supplementary Figure S12.** Uncropped western blots shown in Figure 2 and Supplementary Figure S4

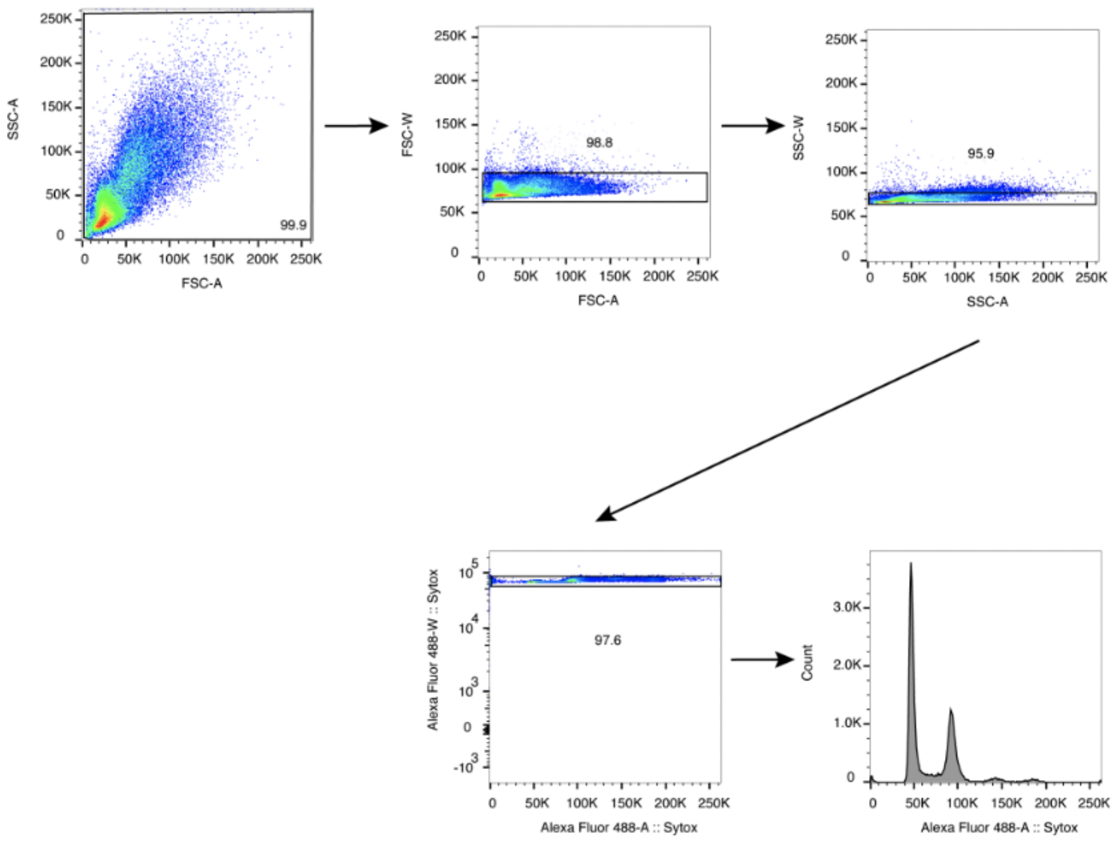


**Supplementary Figure S13.** Gating FACS strategy used in Figure 1D. Shown are wild-type cells 240 minutes after the release from alpha factor block.

**Supplementary Table 1. *Saccharomyces cerevisiae* strains used in this work.**

| Name | Strain | Genotype | Genetic background | Source |
|---|---|---|---|---|
| YMM1 | *wild type (WT)* | *MATa ura3-52 his3Δ200 leu2 lys2-801 ade2-101 trp1Δ63* | *S288c* | |
| YMM5088 | *wild type (WT)* | *MATa his3 leu2 met15 ura3* | *BY4741* | |
| YMM5737 | *gcn5Δ* | *MATa ura3-52 his3Δ200 leu2 lys2-801 ade2-101 trp1Δ63 gcn5Δ::kanMX6* | *S288c* | This study |
| YMM5671 | *esa1-ts* | *MATa ura3-52 his3Δ200 leu2 lys2-801 ade2-101 trp1Δ63 esa1-L254P::KANMX* | *S288c* | This study |
| YMM5686 | *gcn5Δ esa1-ts* | *MATa ura3-52 his3Δ200 leu2 lys2-801 ade2-101 trp1Δ63 esa1-L254P::KANMX gcn5Δ::kanMX6* | *S288c* | This study |
| YMM2936 | *HOS3-NLS-GFP* | *MATa ura3-52 his3Δ200 leu2 ls2-801 ade2-101 trp1Δ63 HOS3-GFP::KAN* | *S288c* | Kumar et al., 2018 |
| YMM3073 | *GAL1pr:HOS3-NLS-GFP MYO1-mCherry* | *MATa ura3-52 his3Δ200 leu2 lys2-801 ade2-101 trp1Δ63 natNT2::GAL1pr-HOS3-NLS-GFP::KAN MYO1-mCherry::hphNT1* | *S288c* | Kumar et al., 2018 |
| YMM5121 | *GAL1pr:HOS3-NLS-GFP* | *ura3-52 his3Δ200 leu2 lys2-801 ade2-101 trp1Δ63 natNT2::GAL1pr-HOS3-NLS-GFP::KAN* | *S288c* | This study |
| YMM5123 | *GAL1pr:HOS3(EN)-NLS-GFP* | *ura3-52 his3Δ200 leu2 lys2-801 ade2-101 trp1Δ63 natNT2::GALpr-hos3-EN(H196E D231A)-NLS-GFP::KAN* | *S288c* | This study |
| YMM3861 | *GAL1pr:HOS3-NLS-GFP MYO1-mCherry ADGEV* | *MATa ura3-52 his3Δ200 leu2 lys2-801 ade2-101 trp1Δ63 natNT2::GAL1pr-HOS3-NLS-GFP::KAN MYO1-mCherry::hphNT1 ADHpr:GAL4-ER-VP16::URA3 (ADGEV)* | *S288c* | Kumar et al., 2018 |
| YMM5761 | *NUP60-GFP* | *MATa ura3-52 his3Δ200 leu2 lys2-801 ade2-101 trp1Δ63 NUP60-GFP::HIS3MX6* | *S288c* | This study |
| YMM5763 | *nup60-KN-GFP* | *MATa ura3-52 his3Δ200 leu2 lys2-801 ade2-101 trp1Δ63 nup60(K467N)-GFP::HIS3MX6* | *S288c* | This study |
| YMM5769 | *esa1-ts NUP60-GFP* | *ura3-52 his3Δ200 leu2 lys2-801 ade2-101 trp1Δ63 NUP60-GFP::HIS3MX6 esa1-L254P::kanMX* | *S288c* | This study |
| YMM5771 | *esa1-ts nup60-KN-GFP* | *ura3-52 his3Δ200 leu2 lys2-801 ade2-101 trp1Δ63 nup60(K467N)-GFP::HIS3MX6 esa1-L254P::kanMX* | *S288c* | This study |
| YMM5027 | *CLN2-HA* | *MATa ura3-52 his3Δ200 leu2 lys2-801 ade2-101 trp1Δ63 CLN2-6xHA::HIS3* | *S288c* | This study |
| JCY2452 | *esa1-ts NUP60-GFP CLN2-HA* | *MATa ura3-52 his3Δ200 leu2 lys2-801 ade2-101 trp1Δ63 NUP60-GFP::HIS3MX6 esa1-L254P::kanMX CLN2-6xHA::hphNT1* | *S288c* | This study |

| | | | | |
|---|---|---|---|---|
| JCY2450 | *esa1-ts nup60-KN-GFP CLN2-HA* | *MATa ura3-52 his3Δ200 leu2 lys2-801 ade2-101 trp1Δ63 nup60(K467N)-GFP::HIS3MX6 esa1-L254P::kanMX CLN2-6xHA::hphNT1* | *S288c* | This study |
| YMM5036 | *mex67-5* | *MATa leu2Δ1 ura3-52 trp1Δ63 mex67-5::natNT2* | *FY86* | Scarcelli et al. 2007 |
| YMM5117 | *SAC3-GFP NUP49-3xmCherry* | *MATa ura3-52 his3Δ200 leu2 lys2-801 ade2-101 trp1Δ63 SAC3-GFP::KAN NUP49-3xmCherry::hphNT1* | *S288c* | This study |
| YMM5119 | *hos3Δ SAC3-GFP NUP49-3xmCherry* | *MATa ura3-52 his3Δ200 leu2 lys2-801 ade2-101 trp1Δ63 SAC3-GFP::KAN NUP49-3xmCherry::hphNT1 hos3Δ::natNT2* | *S288c* | This study |
| YMM5351 | *SAC3-GFP NUP49-3xmCherry* | *MATα ura3-52 his3Δ200 leu2 lys2-801 ade2-101 trp1Δ63 SAC3-GFP::TRP NUP49-3xmCherry::hphNT1* | *S288c* | This study |
| YMM5353 | *SAC3-GFP nup60-KN NUP49-3xmCherry* | *ura3-52 his3Δ200 leu2 lys2-801 ade2-101 trp1Δ63 SAC3-GFP::TRP NUP49-3xmCherry::hphNT1 nup60(K467N)* | *S288c* | This study |
| YMM5675 | *SAC3-GFP NUP49-3xmCherry esa1-ts* | *MATα ura3-52 his3Δ200 leu2 lys2-801 ade2-101 trp1Δ63 SAC3-GFP::TRP1 NUP49-3xmCherry::hphNT1 esa1-L254P::kanMX* | *S288c* | This study |
| YMM5549 | *GAL1pr:sfGFP-CLN2PEST nup60-mCherry* | *MATa his3 leu2 met15 ura3 GAL1pr:sfGFP-CLN2PEST::KAN NUP60-mCherry::hphNT1* | *BY4741* | This study |
| YMM5622 | *hos3Δ GAL1pr:sfGFP-CLN2PEST nup60-mCherry* | *MATa his3 leu2 met15 ura3 GAL1pr:sfGFP-CLN2PEST::KAN hos3Δ::natNT2 NUP60-mCherry::hphNT1* | *BY4741* | This study |
| YMM5557 | *GAL1pr:sfGFP-CLN2PEST nup60-KN-mCherry* | *MATa his3 leu2 met15 ura3 GAL1pr:sfGFP-CLN2PEST::KAN nup60(K467N)-mCherry::hphNT1* | *BY4741* | This study |
| YMM5721 | *hos3Δ GAL1pr:sfGFP-CLN2PEST nupP60-KN-mCherry* | *MATa his3 leu2 met15 ura3 GAL1pr:sfGFP-CLN2PEST::KAN hos3Δ::natNT2 nup60(K467N)-mCherry::hphNT1* | *BY4741* | This study |
| YMM5653 | *nup60-mCherry-FKBP SAC3-FRB-GFP* | *MATα his3Δ1 leu2Δ0 ura3Δ0 LYS+, Can1::Ste2pr-Leu2, Lyp1::, tor1-1, Fpr1::Ura NUP60-mCherry-FKBP::natNT2 SAC3-FRB-GFP::KAN* | *BY4742* | This study |
| YMM5637 | *GAL1pr:sfGFP-CLN2PEST NUP60-mCherry-FKBP* | *MATα his3Δ1 leu2Δ0 ura3Δ0 LYS+, Can1::Ste2pr-Leu2, Lyp1::, tor1-1, Fpr1::Ura NUP60-mCherry-FKBP::natNT2 GAL1pr:sfGFP-CLN2PEST::KAN* | *BY4742* | This study |
| YMM5657 | *GAL1pr:sfGFP-CLN2PEST NUP60-mCherry-FKBP SAC3-FRB* | *MATα his3Δ1 leu2Δ0 ura3Δ0 LYS+, Can1::Ste2pr-Leu2, Lyp1::, tor1-1, Fpr1::Ura NUP60-mCherry-FKBP::natNT2 GAL1pr:sfGFP-CLN2PEST::KAN SAC3-FRB::hphNT1* | *BY4742* | This study |

| | | | | |
|---|---|---|---|---|
| YMM5844 | *SAC3-mCherry-FKBP NUP60-FRB WHI5-GFP esa1-ts* | *MATα his3Δ1 leu2Δ0 ura3Δ0 LYS+, Can1::Ste2pr-Leu2, Lyp1::, tor1-1, Fpr1::Ura SAC3-mCherry-FKBP::natNT2 NUP60-FRB::hphNT1 esa1-L254P::kanMX WHI5-GFP::HIS3MX6* | *BY4742* | This study |
| YMM5848 | *SAC3-mCherry-FKBP NUP60-FRB WHI5-GFP* | *MATα his3Δ1 leu2Δ0 ura3Δ0 LYS+, Can1::Ste2pr-Leu2, Lyp1::, tor1-1, Fpr1::Ura SAC3-mCherry-FKBP::natNT2NT2 NUP60-FRB::hphNT1 WHI5-GFP::HIS3MX6* | *BY4742* | This study |
| YMM5850 | *NUP60-GFP WHI5-mCherry* | *MATa ura3-52 his3Δ200 leu2 lys2-801 ade2-101 trp1Δ63 NUP60-GFP::HIS3MX6 WHI5-mCherry::hphNT1* | *S288c* | This study |
| YMM5854 | *NUP60-GFP WHI5-mCherry esa1-ts* | *MATa ura3-52 his3Δ200 leu2 lys2-801 ade2-101 trp1Δ63 esa1-L254P::kanMX NUP60-GFP::HIS3MX6 WHI5-mCherry::hphNT1* | *S288c* | This study |
| YMM5860 | *nup60-KN-GFP WHI5-mCherry esa1-ts* | *MATa ura3-52 his3Δ200 leu2 lys2-801 ade2-101 trp1Δ63 nup60(K467N)-GFP::HIS3MX6 WHI5-mCherry::hphNT1 esa1-L254P::kanMX* | *S288c* | This study |
| YMM5995 | *hat1Δ NUP60-GFP* | *MATa ura3-52 his3Δ200 leu2 lys2-801 ade2-101 trp1Δ63 NUP60-GFP::HIS3MX6 hat1Δ::natNT2* | *S288c* | This study |
| YMM5998 | *esa1-ts hat1Δ NUP60-GFP* | *ura3-52 his3Δ200 leu2 lys2-801 ade2-101 trp1Δ63 NUP60-GFP::HIS3MX6 esa1-L254P::kanMX hat1Δ::natNT2* | *S288c* | This study |
| YMM5773 | *gcn5Δ esa1-ts NUP60-GFP* | *MATa ura3-52 his3Δ200 leu2 lys2-801 ade2-101 trp1Δ63 NUP60-GFP::HIS3MX6 esa1-L254P::kanMX gcn5Δ::kanMX6* | *S288c* | This study |
| YMM5775 | *gcn5Δ esa1-ts nup60-KN-GFP* | *MATa ura3-52 his3Δ200 leu2 lys2-801 ade2-101 trp1Δ63 nup60-KN-GFP::HIS3MX6 esa1-L254P::kanMX gcn5Δ::kanMX6* | *S288c* | This study |
